# Multiscale Spatial Transcriptomic Atlas of Human Basal Ganglia Cell-Type and Cellular Community Organization

**DOI:** 10.64898/2025.12.02.691876

**Authors:** Bereket Tesfay Berackey, Zhiqun Tan, Ginny Wu, Sujan C. Das, Ren Li, Benjamin Esser, Qiao Ye, Mahsa Nafisi, Samuel S. Park, Pedro Adolfo Sequeira Mendieta, Jillian Berry, Firoza Mamdani, Quan Zhu, Todd C. Holmes, Daofeng Li, Ting Wang, M. Margarita Behrens, Bing Ren, Joseph R. Ecker, Bogdan Bintu, Xiangmin Xu

## Abstract

We generated a multi-region, subcellular-resolution spatial transcriptomic atlas of the human basal ganglia by integrating MERFISH+ and Stereo-seq across four neurotypical donors. These datasets profiled ∼7 million cells spanning the caudate, putamen, nucleus accumbens, and globus pallidus, resolving 60 transcriptionally distinct cell types. We show region-selective, molecular and spatial diversification of medium-spiny-neuron cell types and multiple non-neuronal populations with distinct molecular identities and spatial localizations. Subcellular RNA localization captures somatic size and projection-inferred signatures that reflect direct and indirect pathway topology. Cellular community analyses reveal the enrichment of sub-clusters of astrocytes and oligodendrocytes at striosome–matrix borders, while primate-expanded interneurons are confined to matrix territories. Cross-species mapping uncovers orthologous striosome–matrix organization and conserved dorsolateral-ventromedial gene expression gradients. This atlas provides a foundational molecular and spatial framework for studying human basal ganglia architecture, offering a multi-centimeter scale resource that links cell types, spatial architecture, and subcellular transcript topography across multiple nuclei.

**Highlights:**

- Our multi-centimeter scale spatial taxonomy identifies the precise locations of 60 neuronal and glial cell types of human basal ganglia.
- MERFISH+ and Stereo-seq platforms map consistent spatial modules that align with classical neuroanatomical nuclei.
- D1D2 hybrid MSNs and primate-expanded interneurons show regional and domain specific organization
- Subcellular RNA localization reports soma morphology and projection-inferred signatures.

## INTRODUCTION

The human basal ganglia is a critical brain structure involved in motor control, learning and emotion. It is comprised of distinct anatomical nuclei / subdivisions including the striatum (caudate nucleus and putamen), the globus pallidus (external and internal segments, GPe and GPi), and the nucleus accumbens (NAc) ^1^. Basal ganglia cell types and architecture are well characterized in rodents and non-human primates, but are much less understood in humans. The literature from both animal models and humans shows that GABAergic medium spiny neurons (MSNs) are the dominant neuronal cell type in the striatum (85–95%) ^2–5^; MSNs are divided into two major populations: D1-type MSNs, which express the dopamine receptor D1 (DRD1), tachykinin precursor 1 (TAC1), and dynorphin; and D2-type MSNs, which express the dopamine receptor D2 (DRD2), proenkephalin (PENK), and enkephalin ^2,3,6^. The D1 and D2 populations are spatially organized into interdigitated striosome and matrix compartments that form the structural foundation of basal ganglia function ^3,7,8^. Further studies, including those using single-nucleus RNA-seq in animal and human brains have revealed additional D1/D2 neuronal diversity ^9,10^, and identified multiple classes of interneurons, including cholinergic (choline acetyltransferase-positive; ChAT+), parvalbumin-positive (PV+), somatostatin-positive (SST+), and neuropeptide Y/tyrosine hydroxylase-positive (NPY+/TH+) subtypes ^10–14^. The nucleus accumbens, located at the ventral extension of the striatum, contains both D1 and D2 MSNs and is further divided into core and shell subregions. These subregions mirror the cellular composition of the dorsal striatum, including diverse interneuron classes ^10,15^. The globus pallidus consists of two principal divisions: the external segment (GPe) and the internal segment (GPi) ^3,16^. Both regions are dominated by PV+ GABAergic projection neurons ^17^. Within GPe, prototypic neurons express PV, LIM homeobox 6 (LHX6), and NK2 homeobox 1 (NKX2-1) ^18,19^; arkypallidal neurons express neuronal PAS domain protein 1 (NPAS1) and forkhead box protein P2 (FOXP2) ^20,21^. GPi contains PV+/LHX6+ projection neurons that form the principal output pathway of the basal ganglia ^16,17^.

A comprehensive understanding of human basal ganglia organization requires integrating cell-type identity, molecular signatures, and spatial architecture within an intact anatomical framework. Traditional single-cell RNA sequencing provides molecular resolution but lacks spatial context, limiting its ability to capture circuit-level organization. Emerging spatial transcriptomic technologies map gene expression directly within intact tissue sections that retain the spatial relationships of cell types and structures ^10,14,22–24^. These methods include probe-based multiplexed fluorescent in situ hybridization (FISH) and next-generation sequencing (NGS)-based spatially resolved transcriptomics, each with complementary strengths in resolution, transcriptome coverage, and throughput ^23,25,26^. These approaches enable an unprecedented mapping of brain architecture through quantitative classification of cell types and detection of spatially patterned gene expression. Although previous studies have revealed the spatial organization of striosomes within the striatal matrix and transcriptional gradients across basal ganglia nuclei in model organisms such as mice and macaques ^10,27^, a comprehensive molecular and cell-type-resolved spatial atlas of the adult human basal ganglia remains lacking. This gap has persisted in part because of the large size and complex cytoarchitecture of human basal ganglia nuclei, as well as technical limitations in the spatial scale and resolution of existing mapping approaches Addressing this limitation is particularly critical given the central role of the basal ganglia in human movement disorders (including Parkinson’s disease and Huntington’s disease), neuropsychiatric conditions, and species-specific differences in basal ganglia cellular composition and structural organization that reflect the complexity of inputs and outputs for these structures in humans ^7,28^.

Recent advances in large-scale spatially resolved transcriptomics have transformed our ability to characterize cellular heterogeneity and molecular architecture within intact neural tissue such as human basal ganglia. Two particularly powerful approaches are Multiplexed Error-Robust Fluorescence In Situ Hybridization (MERFISH) ^26,29–33^ and Spatio-Temporal Enhanced Resolution Omics-sequencing (Stereo-seq) ^34–38^. MERFISH enables highly multiplexed, single-molecule RNA imaging at subcellular resolution, allowing the precise mapping of hundreds to thousands of genes within individual cells while maintaining their spatial context ^26,29,31^. This approach has been instrumental in decoding cell-type organization across cortical and subcortical structures in multiple species that reveal both conserved and species-specific architectural principles in the mouse, monkey and human brains ^14,27,36,39–41^. Subsequent advancements in MERFISH have expanded high-resolution spatial transcriptomics into a multimodal platform capable of imaging DNA, RNA, and proteins within the same tissue specimen ^42,43^. Most recently, the incorporation of modified probe designs, high-throughput microscopy, and microfluidic automation has led to MERFISH+ ^44^, an enhanced implementation that enables imaging across multi-centimeter–scale tissue areas. This extension achieves large-area spatial transcriptomics with subcellular precision for integrative molecular and structural analyses ^44^. Stereo-seq employs DNA nanoball-patterned arrays to generate spatially barcoded cDNA libraries that capture transcriptomes from large tissue sections at high resolution ^34,36–38^. Stereo-seq has been successfully applied to generate whole-brain atlases in mice and to characterize cortical and subcortical organization in nonhuman primates ^34–38^. This sequencing-based approach provides an unbiased, genome-wide view of spatial gene-expression patterns across extensive anatomical domains, albeit at a lower spatial resolution compared with imaging-based methods such as MERFISH+.

In this study, we harness complementary strengths of MERFISH+ and Stereo-seq to generate a multiscale spatial transcriptomic atlas of the human basal ganglia to capture spatial gene-expression patterns, specific cell types, and molecular architectures across multi-centimeter tissue sections from four neurotypical adult postmortem donors. This atlas offers a novel view of human basal ganglia organization of cell types, cellular communities, and region-specific transcriptional gradients at subcellular precision that was previously uncharted.

## RESULTS

### Multi-region, subcellular-resolution spatial transcriptomic mapping of human basal ganglia

We obtained four postmortem human brain fresh frozen slabs across anterior-to-posterior segments of the basal ganglia encompassing the caudate nucleus, putamen, nucleus accumbens (NAc), globus pallidus, and internal capsule (Figure 1A). Tissue samples were collected from four neurotypical Caucasian male donors, aged 36–53 years, with no pathological evidence of neurodegenerative disease or other brain injury. RNA quality was confirmed by RNA Integrity Number (RIN) values ranging from 7.0 to 8.4, indicating excellent preservation for downstream analyses (Table S1).

**Figure 1.**
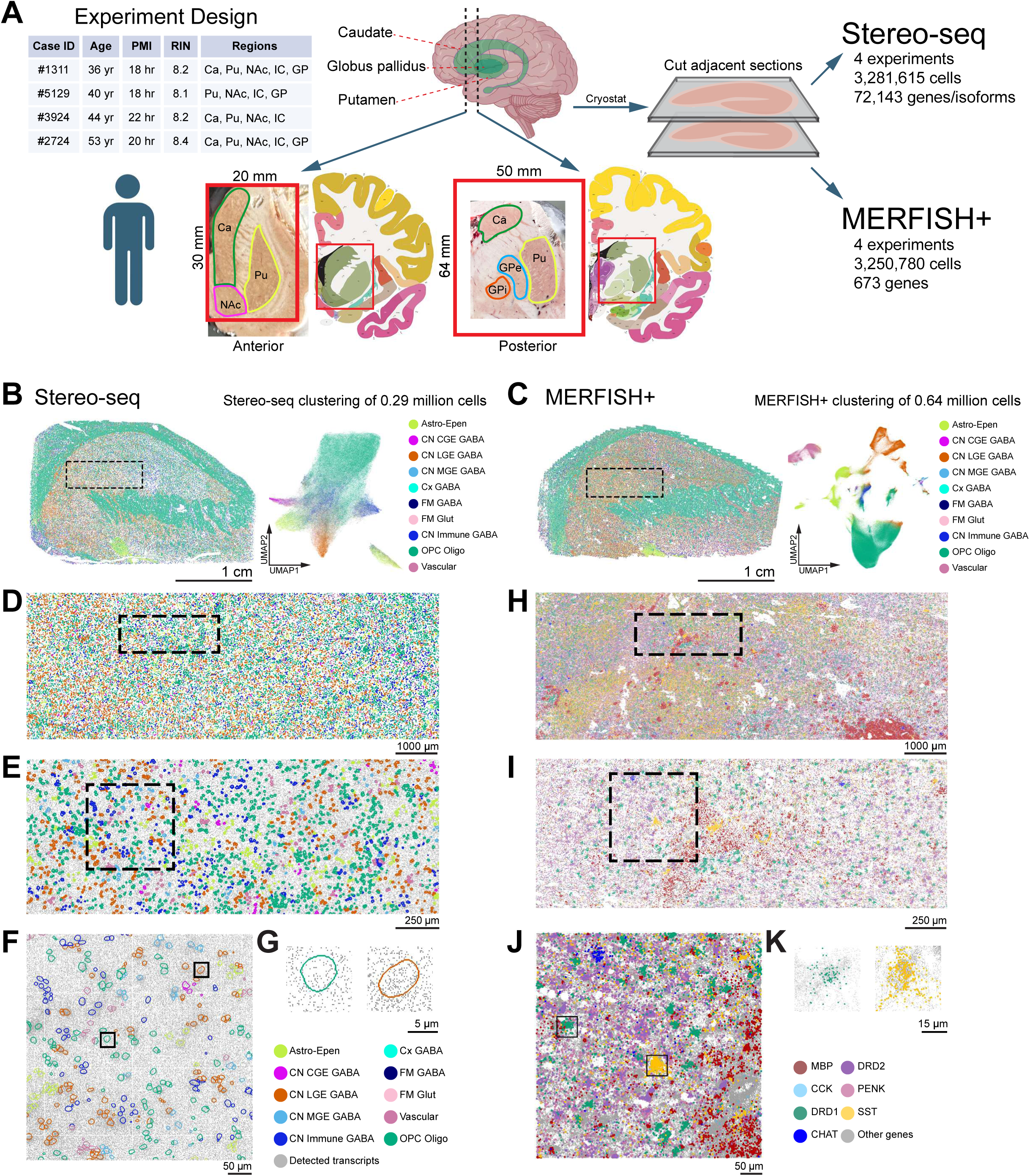
Multi-centimeter, subcellular-resolution spatial transcriptomic mapping of the human basal ganglia with Stereo-seq and MERFISH+. (**A**) Overview of study design, donor and sample quality and metadata, tissue sampling strategy, and adjacent-section workflows. (**B**) Stereo-seq dataset from an anterior basal ganglia section (donor case ID 2724): left, spatial map with cell-type assignments; right, UMAP embedding resolving 10 major cell classes from the same section. (**C**) MERFISH+ dataset from an adjacent anterior section. Left, spatial map with cell-type assignments; right, UMAP embedding resolving 10 major cell classes corresponding to Stereo-seq in (**B**). (**D–G**, progressively greater magnification) Multi-scale Stereo-seq enlarged views of the boxed region in (**B**), illustrating the resolution span from centimeter-scale tissue anatomy to micron-scale detection of individual transcripts, with corresponding cell-type labels (**D, E**), cell-contour segmentation (**F**), and single-cell detail (**G**). (**H–K**, progressively greater magnification) Multi-scale MERFISH+ enlarged views of the boxed region in (**C**), showing transcripts for selected genes (DRD1, DRD2, PENK, CCK, SST, CHAT, MBP) (color coded) and other genes (gray coded) that define cell-types at micron-scale resolution. See color keys at (**K**). Scale bars as indicated

For spatial transcriptomic profiling, 10 µm-thick cryostat sections were processed for Stereo-seq with the coverage area size of up to 20 mm × 30 mm, while adjacent 16 µm-thick sections were selected for MERFISH+ imaging with an area size of up to 43 mm × 30 mm (Figure 1A, Supplementary Movie 1, Table 1S & Figure 1S in Supplemental Information). A single 20mm x 30mm Stereo-seq chip captured ∼0.9 million cells (with ∼0.29 million cells selected for cell type classification), which were clustered into 10 different major cell types (Figure 1B). Across four Stereo-seq assays, we obtained genome-wide transcriptomic coverage of ∼70,000 genes across ∼3.2 million cells. In parallel, comparable numbers of cells per area were captured by MERFISH+ on adjacent sections targeting a panel of 673 selected genes (Figure 1C, Supplemental Data). Four MERFISH+ experiments captured ∼3.9 million cells at single-molecule resolution over multi-centimeter-scale tissue areas. Both methods yielded highly concordant results for the spatial distributions of major cell types across corresponding anatomical regions (Figure S1A). Although Stereo-seq detects fewer transcripts per cell due to its sequencing-based capture efficiency, the expression levels of the 673 MERFISH+ target genes show strong cross-platform correlation between the two modalities (Pearson correlation coefficient of 0.71), demonstrating the quantitative agreement of spatial gene-expression measurements across methodological approaches (Figure S1B) with high reproducibility across and between experiments (Figure S1C-D). Although Stereo-seq provides transcriptome-wide coverage, its spatial signal localization is inherently more diffuse (Figure 1D-G) than MERFISH+ that delineates sharp cellular boundaries based on transcript density (Figures 1H–K; Supplementary Movie 1). Given these respective advantages and limitations, we used MERFISH+ to focus on fine-scale cell definition and subcellular localization, and Stereo-seq for exploring large scale gene expression patterns across centimeter scale tissue.

### Molecular and spatial taxonomy of cell types in the human basal ganglia

Using MERFISH+, we resolve approximately 60 transcriptionally distinct cell types that are separable when embedded in Uniform Manifold Approximation and Projection (UMAP) space of gene-expression data (Figure 2A). These cell types show expression of expected corresponding marker genes (Figure S2) and demonstrate strong correspondence with single-nucleus RNA sequencing (snRNA-seq) datasets from the Allen Institute Mammalian Basal Ganglia Consensus Cell Type Atlas (Allen Institute; RRID:SCR_024672) ^45^ (Figure 2B), validating our cell type classification. We identify twelve major classes spanning anterior-to-posterior sections (Figure 2C). Among these are eight neuronal classes that include caudal and lateral ganglionic eminence (CGE/LGE)-derived GABAergic neurons, which are dominated by medium spiny neurons (MSNs) (Figure S3A).

**Figure 2.**
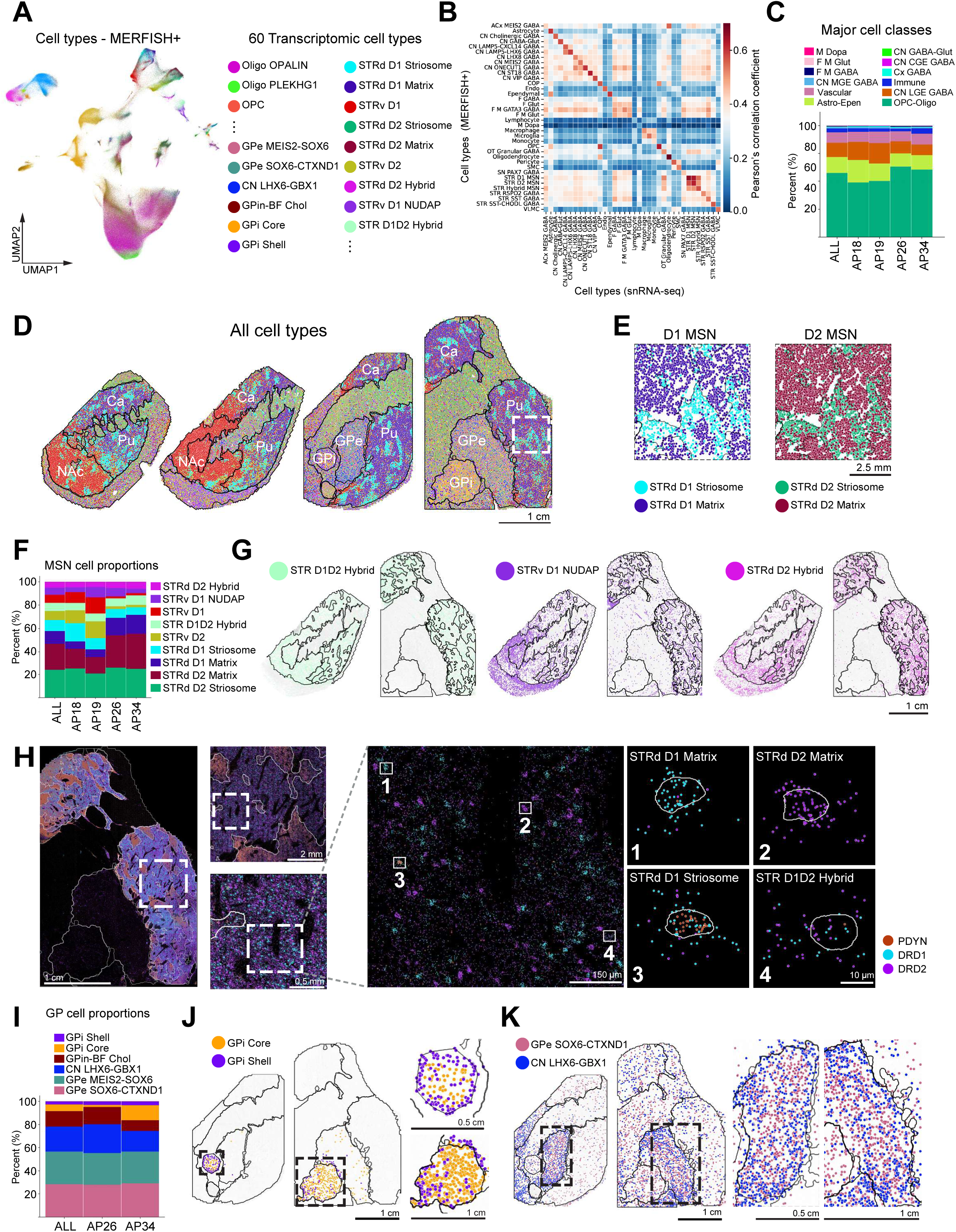
Spatial organization of major neuronal cell types across the human basal ganglia. (**A**) UMAP embedding of MERFISH+ single cell data with 60 transcriptionally defined cell types classified using MapMyCell (Allen Institute; RRID:SCR_024672; https://github.com/AllenInstitute/cell_type_mapper). (**B**) High correlation between MERFISH+ defined cell types and snRNA-seq–defined clusters. (**C**) Fractional composition of 12 major cell types for all sections combined and across the anterior–posterior (A–P) axis / position (AP 18a, 19a, 26a, 34a), defined based on Allen institute human brain atlas^59^ (stacked bars). (**D**) Spatial distributions of MERFISH+ mapped cell types in the brain section samples across the AP axis (left - right AP 18a, 19a, 26a, 34a). See cell type color keys in (**A**). The contour lines in the sample sections / maps (i.e., panels **D**, **G, H** and **J-K**) shown in this figure and subsequent figures denote the anatomical regional boundaries derived from the cellular community analysis (further described later in the results section of “Mesoscale cellular community architecture of the human basal ganglia”). (**E**) Zoomed-in view of D1 and D2 MSNs within the boxed region of the posterior section (AP 34a) in (**D**). (**F**) Fractional composition of MSN subtypes for all sections combined, then along the A–P axis (AP 18a, 19a, 26a, 34a shown for each section stacked bars). (**G**) Spatial distributions of STR D1D2 Hybrid, STRv D1 NUDAP, and STRd D2 Hybrid MSNs at anterior (AP 19a) and posterior (AP 34a) sections. (**H**) Multiscale spatial gene-expression maps for PDYN, DRD1, and DRD2 in a posterior section (AP 34a) showing PDYN-enriched striosomes and D1/D2 MSNs predominantly in the putamen matrix territories. Boxed regions show sequential progressively magnified centimeter→millimeter→micron-scale views. Cell-segmented higher-magnification panels highlight representative gene expression of DRD1 in D1 MSN (1), DRD2 in D2 MSN (2), DRD1 and PDYN in STRd D1 Striosome (3), and both DRD1 and DRD2 in STR D1D2 Hybrid (4) cells. (**I**) Fractional composition of globus pallidus (GP) neuronal subtypes for all combined sections and for AP 26a and AP 34a (stacked bars). (**J**) Spatial distributions of GPi Core and GPi Shell neurons, with enlarged views illustrating the anatomical topographies of each subtype at AP 26a and AP 34a. (**K**) Spatial distributions of GPe SOX6-CTXND1 and CN LHX6–GBX1 neuronal subtypes, with enlarged views of boxed regions in GPe at AP 26a and AP 34a.

We further examined MSNs – the predominant neuronal population of the striatum that are essential for modulating movement control in the basal ganglia circuit ^3^. MSNs segregate into two canonical classes – D1 and D2 MSNs – further subdivided into six major dopaminergic MSN subtypes: STRd D1 Striosome, STRd D1 Matrix, STRv D1, STRd D2 Striosome, STRd D2 Matrix and STRv D2 (dorsal striatum, STRd; ventral striatum, STRv) (Figure 2D-E, Figure S3B). These subtypes display distinct dorsal–ventral and striosome–matrix distribution patterns across basal ganglia compartments along the anterior-posterior axis (Figure 2D, Figure S3B). Specifically, D1 and D2 striosomal MSNs are enriched within striosome compartments, whereas D1 and D2 matrix MSNs predominate in the surrounding matrix (Figure 2E).

The proportion of MSN cell types is largely consistent along the anterior-posterior axis (Figure 2F). D2 MSNs constitute the largest proportion compared to D1 MSNs (approximately 54 % vs 27 %, respectively). D1 and D2 MSNs together represent ∼83 % of all striatal neurons.

Beyond these canonical MSN populations, our analyses uncover spatial distributions of several novel and rare MSN cell types described across species including mouse and non-human primate that are “mixed-identity” neurons that co-express D1- and D2-associated gene modules – these neurons are not regionally restricted cell types to discrete striatal or pallidal territories ^46–48^. The D1–D2 hybrid (STR D1D2 Hybrid), striosome-matrix hybrid (STRd D2 Hybrid), and D1 NUDAP (Neurochemically Unique Domains in the Accumbens and Putamen) (STRv D1 NUDAP) MSNs are primarily localized within the putamen, caudate nucleus, and nucleus accumbens (Figure 2G), but are largely absent from the globus pallidus and internal capsule. The mixed state of these cell populations is corroborated by colocalization within individual neurons of multiple transcripts of DRD1, DRD2, and prodynorphin (PDYN), a canonical striosomal MSN marker (Figure 2H).

We next examined neuronal subtypes within the globus pallidus. Analysis of two posterior sections encompassing both internal (GPi) and external (GPe) segments reveal six transcriptionally distinct GP-associated neuronal classes: GPi Shell, GPi Core, GPin-BF Cholinergic, CN LHX6–GBX1, GPe SOX6–CTXND1, and GPe MEIS2–SOX6 (Figure 2I, Figure S4). As indicated by their nomenclature, GPi Shell and GPi Core neurons are localized to peripheral and central GP regions, respectively (Figure 2J). In contrast, GPe SOX6–CTXND1 and CN LHX6–GBX1 neurons are concentrated within the GPe and show only sparse distributions in other basal ganglia nuclei (Figure 2K).

In addition to neuronal classes, we uncovered multiple non-neuronal populations with distinct molecular identities and spatial localizations (Figure 3A, Figure S5). Four major categories are observed: immune, vascular, astrocytic/ependymal (Astro–Epen), and oligodendrocyte lineage (OPC–Oligo) cells, each exhibiting characteristic regional patterns. OPC–Oligo cells represent the oligodendrocyte lineage and form the most abundant non-neuronal population, comprising 60–70% of all non-neuronal cells (Figure 3A-left). This group includes five transcriptionally distinct subtypes: committed oligodendrocyte progenitors (COP), oligodendrocyte progenitor cells (OPC), Oligo PLEKHG1, immune-associated oligodendrocytes (ImOligo), and Oligo OPALIN (Figure 3A-right). Notably, Oligo OPALIN and Oligo PLEKHG1 display complementary spatial distributions (Figure 3B) with differential transcriptional signatures (Figure 3C). We found that Oligo OPALIN cells are enriched along striatopallidal fiber tracts (arrows, Figure 3B-right), whereas Oligo PLEKHG1 cells are preferentially localized to the internal capsule white matter and globus pallidus, but are sparse in the putamen and caudate nucleus — this pattern is consistent across different sections (Figure S6A).

**Figure 3.**
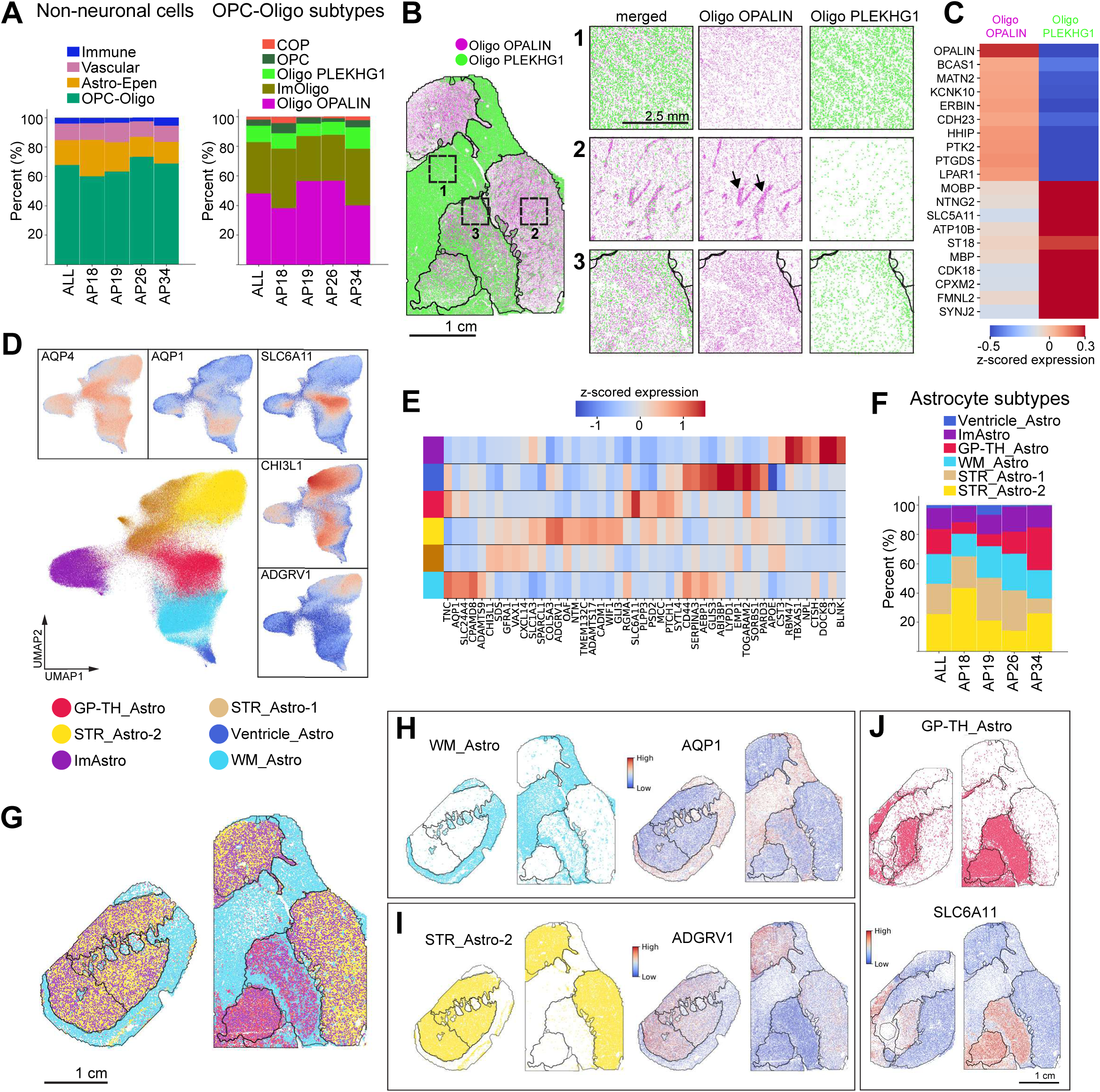
Spatial organization of major non-neuronal cell types across the human basal ganglia. (**A**) Fractional composition of major non-neuronal classes (left) and OPC/oligodendrocyte (Oligo) subtypes (right) for all combined sections and across the anterior–posterior axis (AP 18a, 19a, 26a, 34a). (**B**) Spatial distributions of two oligodendrocyte subtypes, Oligo-OPALIN and Oligo-PLEKHG1. Zoomed-in views of the boxed regions: (1) internal capsule, (2) putamen with striatopallidal fiber bundles (arrows), and (3) GPe. (**C**) Heatmap (log1p z-score) of the top differentially expressed genes distinguishing Oligo-OPALIN and Oligo-PLEKHG1 subtypes. (**D**) UMAP of six subtypes of astrocytes defined based on the MERFISH+ astrocytic genes measured, with expression maps for representative differentially expressed astrocytic genes. (**E**) Heatmap (log1p z-score) of the top differentially expressed genes associated with each astrocyte subtype. (**F**) Fractional composition of astrocyte subtypes for all combined sections and across the anterior–posterior (AP) axis (left - right: AP 18a, 19a, 26a, 34a, stacked bars). (**G-J**) Spatial distributions of the six astrocyte subtypes in anterior (AP 18a) and posterior (AP 34a) sections (**G**), including (**H**) white-matter–enriched astrocytes (WM_Astro) localized to the internal capsule with high AQP1 expression (right), (**I**) striatum-enriched astrocytes (left) with high ADGRV1 expression (right), and (**J**) globus pallidus-enriched astrocytes (GP-TH_Astro) with high SLC6A11 expression. Color keys are consistent across panels (**G–J**). Astrocyte cell type key as shown in (**D**). Scale bars as indicated.

We found that astrocytes also exhibit pronounced regional heterogeneity. Subclustering analysis identifies six astrocyte subtypes, each defined by distinct marker gene combinations that include aquaporin 4 (AQP4), aquaporin 1 (AQP1), solute carrier family 6 member 11 (SLC6A11), chitinase 3-like 1 (CHI3L1), and adhesion G protein-coupled receptor V1 (ADGRV1) (Figure 3D). These subtypes form discrete clusters in UMAP space with divergent transcriptional signatures (Figure 3E). Except for the ventricle-associated astrocytes (Ventricle_Astro, captured in section AP19), the relative abundances of astrocyte subtypes are generally consistent across anterior and posterior basal ganglia regions (Figure 3F). Spatial mapping reveal distinct topographic specializations (Figure 3G and Figure S6B-C): WM_Astro cells, characterized by high AQP1 expression, are localized primarily to white matter tracts and the peripheral globus pallidus (Figure 3H); STR_Astro-2 cells, defined by ADGRV1, are enriched in striatum (Figure 3I), and GP-TH_Astro cells, expressing SLC6A11, are selectively concentrated in the globus pallidus (Figure 3J). The differential gene expression of these varied astrocytic cell types likely provides clues for their specific function. For instance, SLC6A11 (also known as GAT-3) is a GABA transporter that is important for GABA uptake and metabolism in astrocytes^49,50^, consistent with the high abundance of GABAergic neurons in the globus pallidus. These findings underscore the molecular, cellular, and spatial heterogeneity of glial and neuronal populations that define the complex architecture of the human basal ganglia.

### Subcellular transcript localization for somatic size estimation and projection inference

The high spatial precision of MERFISH+ enables the localization of individual RNA molecules at subcellular resolution (Figure 1J-K; Supplementary Movie 1). This allows each transcript to be categorized as nuclear (within the nucleus), somatic (within the cytoplasm), or putative cellular processes (distributed along processes or extensions) (Figure 4A) (see STAR Methods). While nuclear and somatic transcripts are used to define cell identity, we developed additional analysis to capture somatic morphology of single neurons. We also used the RNA distributions in putative cellular processes to reveal information about projection topology in human basal ganglia.

**Figure 4.**
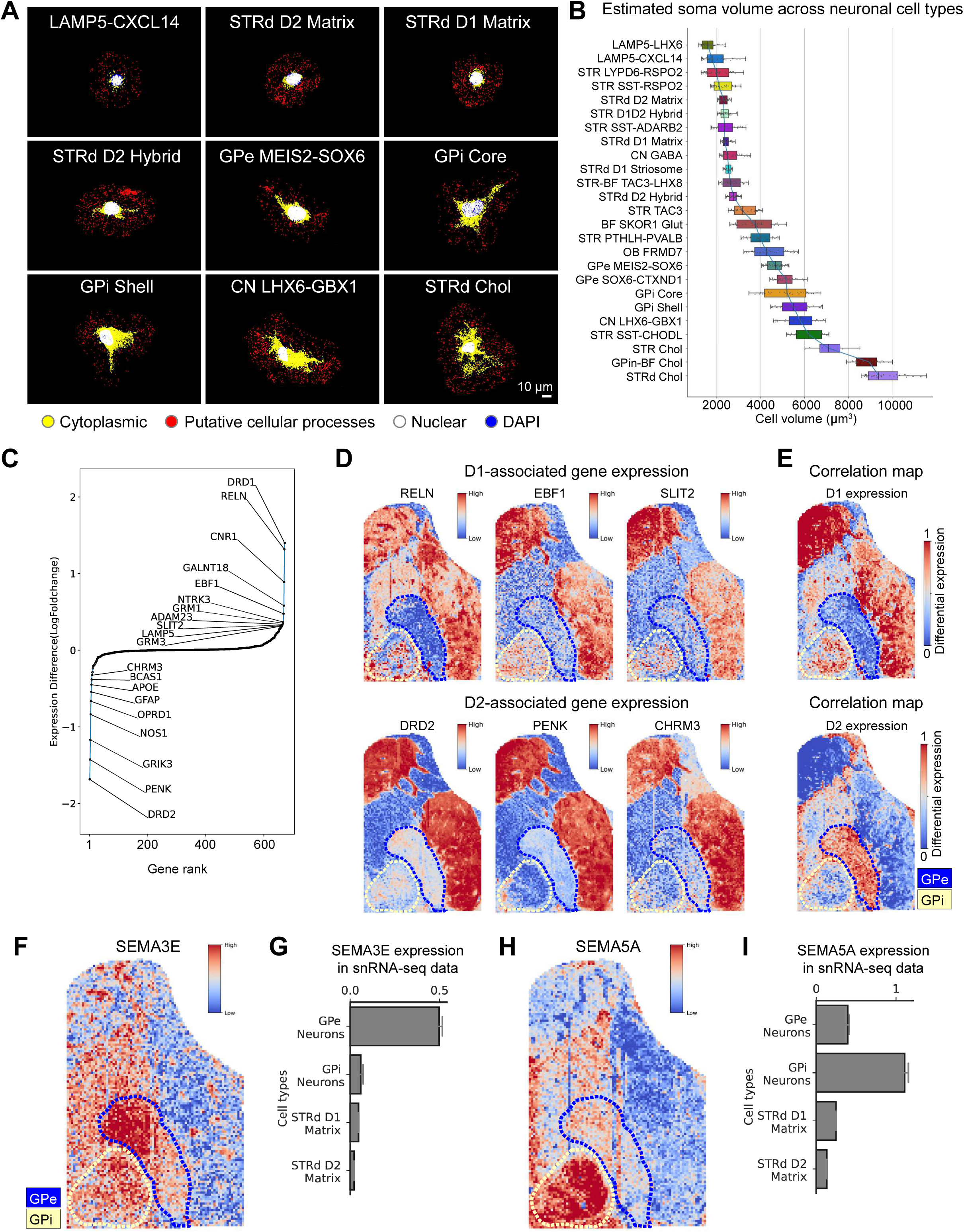
Subcellular MERFISH+ transcript localization infers cell-type-specific somatic morphology and projection signatures. (**A**) Images of MERFISH+ transcripts pseudo-colored to indicate subcellular localization: nuclear (white), cytoplasmic (yellow), and putative cellular processes (red) across nine selected cell types. Distinct localization profiles highlight cell-type–specific soma size, shape, and putative neuritic extent. DAPI shown in blue but largely masked by the nuclear transcript signal. (**B**) Estimated soma means and ranges of volumes (µm³) inferred from transcript-density segmentation across neuronal cell types in the human basal ganglia, including those shown in (**A**). (**C)** Plot of differential gene expression between D1 and D2 matrix MSNs measured by MERFISH+. Genes are ranked based on differential expressions. Top differential expressed genes are annotated. **(D)** Spatial maps of transcripts localized in putative cellular processes for genes enriched in D1 matrix MSNs (RELN, EBF1, SLIT2; top) and D2 matrix MSNs (DRD2, PENK, CHRM3; bottom). (**E**) Section-wide correlation maps of D1 matrix MSN (top) and D2 matrix MSN (bottom) gene expression with the expression in putative cellular processes across the entire posterior section (AP34a). Dashed blue outline marks GPe and dashed yellow outline marks GPi. Projection-biased localization reflects direct (D1→GPi) versus indirect (D2→GPe) pathway topology. **(F)** Gene expression map of SEMA3E transcripts localized in putative cellular processes in the posterior section (AP34a). **(G)** The log1p normalized count of nuclear SEMA3E transcripts obtained from snRNA-seq data. **(H)** Gene expression maps of SEMA5A transcripts localized in putative cellular processes in the posterior section (AP34a). **(I)** The log1p normalized count of nuclear SEMA5A transcripts obtained from snRNA-seq data.

To estimate somatic size, we modeled each cell’s cytoplasmic space as a three-dimensional point cloud derived from somatic RNA coordinates, which was then converted into a volumetric mesh representation (see STAR Methods). This analysis reveals substantial heterogeneity in neuronal soma volumes across basal ganglia regions (Figure 4A-B). Cholinergic interneurons and globus pallidus neurons including GPi Core, GPi Shell, CN LHX6–GBX1, and GPe MEIS2–SOX6 populations, exhibit the largest median soma volumes (up to ∼9,400 µm³), whereas LAMP5+ interneurons and MSNs display markedly smaller volumes (∼1,800 µm³ and ∼2,500 µm³, respectively) (Figure 4B). This is consistent with histological studies in mouse and human basal ganglia ^51–53^. Although MSN subtypes are more consistent in soma volumes, other neuronal classes exhibit more subtype-dependent variability; for example, SST–CHODL interneurons have among the largest median volumes (∼6,200 µm³), while SST–ADARB2 interneurons are significantly smaller (∼2,300 µm³) (p-value = 6.9 x 10^-15^, Wilcoxon rank-sum test). These quantitative reconstructions highlight the pronounced morphological diversity among neuronal subtypes in the human basal ganglia.

We next analyzed RNA distributions in putative cellular processes including axonal and dendritic processes (“neurite transcripts”) which provide indirect information about neuronal projections. Focusing on MSNs, in the mouse and monkey basal ganglia circuitry, D1 matrix MSNs project directly to the GPi and substantia nigra pars reticulata, whereas D2 matrix MSNs form the indirect pathway via the GPe ^3,54^. Consistent with this organization, we found that D1-enriched transcripts (e.g., RELN, EBF1, SLIT1) exhibit higher extra-somatic signal density within the GPi, while D2-enriched transcripts show preferential accumulation in the GPe (Figure 4C–D). Quantitative correlation analysis across the 673 genes imaged with MERFISH+ confirms this relationship: D1-specific genes display significantly higher spatial correlation scores in the GPi than in the GPe, whereas the inverse pattern is observed for D2-specific genes (Figure 4E). In addition, we observed that GPi and GPe regions have high concentration of extrasomatic SEMA3E and SEMA5A, ligands part of the semaphorin family of axonal guidance molecules ^55,56^. We found SEMA3E is enriched in GPe (Figure 4F), while SEMA5A is prominently found in GPi (Figure 4H). Their expression is also higher in the nuclei as confirmed using the snRNA-seq data from the Allen Institute Basal Ganglia Consensus Cell Type Atlas (Figure 4G, I). This might reflect that specific semaphorin gene family members mediate long range connections in neurons such as MSNs targeting specific segments of the globus pallidus ^55,57^. Together, these findings demonstrate that MERFISH+ not only resolves molecular and spatial identities of cells but also captures morphological and projection-inferred features encoded in the subcellular transcriptome and provides an unprecedented window into the structural and functional organization of human basal ganglia circuits.

### Mesoscale cellular community architecture of the human basal ganglia

To resolve mesoscale organization beyond individual cell types, we performed cellular community analysis ^14,58^, which groups cells based on the composition of their spatial neighborhoods. Specifically, for each cell, we calculated the frequency of neighboring cell types within a radius of 250 - 400 µm and then constructed neighborhood composition vectors which were subsequently clustered into spatial modules (STAR Methods and Figure 5A). This approach yields 14 discrete spatial modules that correspond to anatomical regions defined in the Allen Human Brain Atlas ^59^ (Figure 5B).

**Figure 5.**
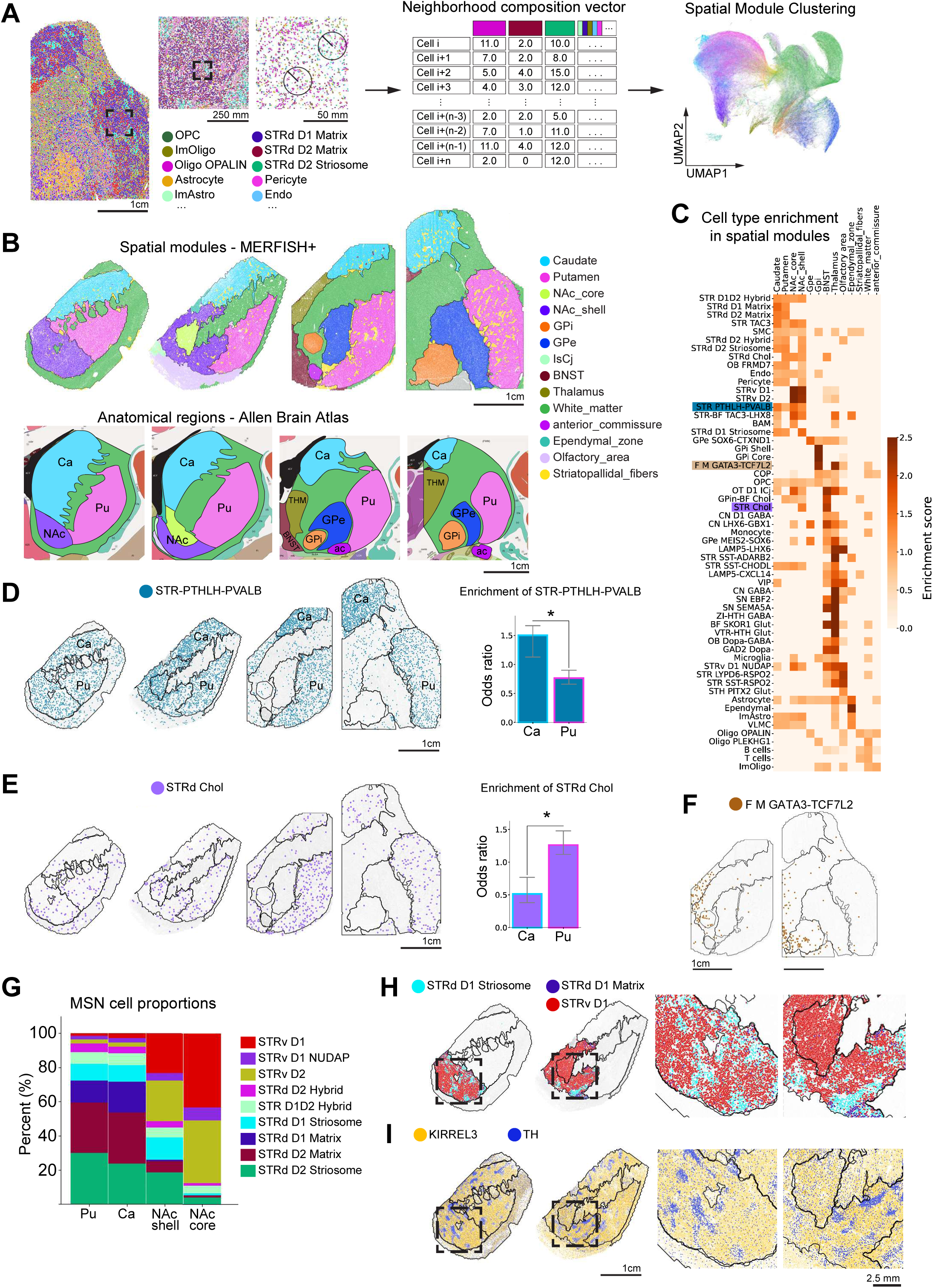
Cellular community architecture recapitulates major anatomical modules of the human basal ganglia. (**A**) Spatial map of major cell types of the posterior section (as shown in Figure 2D) and inferred cellular community modules derived from the local neighborhood composition analysis, accompanied by a UMAP embedding of community-level clustering across major cell classes. (**B**) Top: Community module assignments mapped onto anatomical regions across the anterior–posterior (A–P) axis (AP 18a, 19a, 26a, 34a), capturing mesoscale organization. Bottom: Corresponding anatomical reference images from the Allen Human Brain Atlas^59^. Pu, putamen; Ca, caudate; NAc, nucleus accumbens; GP, globus pallidus. Color keys are consistent across panels in (**B**). (**C**) Heatmap showing cell-type enrichment scores for each community module (rows = cell types; columns = modules). (**D**) Spatial distribution of fast-spiking interneurons (STR FS PTHLH–PVALB) across A–P levels, highlighting preferential enrichment in caudate. Right: odds-ratio plot quantifying caudate > putamen representation (* p = 0.021; Wilcoxon rank-sum). (**E**) Spatial distribution of cholinergic interneurons (STRd ChAT+) across the A–P axis, showing relative enrichment in putamen compared to caudate. Right: odds-ratio plot quantifying putamen > caudate representation (* p = 0.021; Wilcoxon rank-sum). (**F**) Spatial map showing the enrichment of the cell type F M GATA3-TCF7L2 in the internal segment of the globus pallidus (GPi). (**G**) Fractional composition of MSN subtypes within putamen (Pu), caudate (Ca), nucleus accumbens (NAc) core, and NAc shell. (**H**) Spatial distributions of D1 MSN subtypes in the NAc from two anterior sections: AP 18a (shell; left zoom-in) and AP 19a (shell + core; right zoom-in), illustrating enrichment of STRd D1 striosome-like MSNs in shell relative to core regions. (**I**) Marker gene maps illustrating compartmentalization of striosome–matrix-like architecture in the NAc shell: tyrosine hydroxylase (TH) (striosome-associated) and KIRREL3 (matrix-associated).

Each spatial module display characteristic cell-type enrichment patterns corresponding to known anatomical domains ^13^. For example, GPe SOX8–CTXND1 and GPe MEIS2–SOX6 neurons are enriched in the external globus pallidus (GPe), whereas GPi Core and GPi Shell neurons are confined to the GPi. Similarly, STRv D1 and STRv D2 MSNs are enriched in the nucleus accumbens shell and core, while STRd D1 and STRd D2 MSNs populate the striatum (caudate nucleus and putamen) (Figure 5C).

Besides these canonical distributions, several interneuron and projection-type populations exhibit distinct regional biases. For instance, STR fast-spiking PTHLH–PVALB GABAergic interneurons are preferentially enriched in the caudate nucleus (Figure 5D, in agreement with previous snRNA-seq analysis ^11^. STRd ChAT+ GABAergic interneurons are more abundant in the putamen (Figure 5E). In contrast, FM GATA3–TCF7L2 neurons, a transcriptionally distinct population with unclear lineage affiliation are enriched in the GPi nucleus of the basal ganglia; ; this represents a previously unknown neuronal subtype within this region (Figure 5F).

Quantitative analysis of MSN subtype proportions across striatal subregions reveals that the caudate nucleus and putamen share comparable D1/D2 ratios (Figure 5C). In contrast, both D1- and D2-striosomal MSNs are detected in the NAc shell but are largely absent from the NAc core (Figure 5G–H). This differential organization is corroborated by the expression of established molecular markers: TH associated with striosome compartments ^3^, is enriched in the NAc shell, whereas KIRREL3, a matrix-associated marker ^60^, is predominantly expressed in the NAc core (Figure 5I).

These findings demonstrate that cellular community analysis reconstructs macroscopic anatomical boundaries and allows for the identification of cell types located within or absent from the anatomical regions comprising the human basal ganglia circuits.

### Striosome–matrix domains and cross-species orthologous organization

Beyond its mesoscale structures, the striatum exhibits micro-domain architecture composed of striosome and matrix compartments ^61^. To delineate these domains, we further applied cellular community analysis restricted to MSN subtype composition—the principal determinant of striosome–matrix identity (Figure 6A-B). Beyond canonical D1/D2 MSN distributions, we identify additional cell types exhibiting previously uncharacterized striosome–matrix enrichments. Notably, STR TAC3–PLPP4 interneurons are exclusively localized to the matrix compartment (Figure 6C, 6D top). These interneurons are primate-specific^62,63^ and likely represent the orthologous lineage of rare Tac2+/TH+ interneurons that is described in rodents ^17,64^. In our human dataset, STR TAC3–PLPP4 neurons are among the most abundant population of striatal interneurons (Figure 6E). This expansion is consistent with the trend of expansion for this cell type seen in prior primate and human studies, whereas their putative mouse homologs represent only ∼6% of interneurons ^62–64^. This marked expansion of TAC3–PLPP4 interneurons in primates suggests an evolutionary adaptation that supports the integration of the greater density of prefrontal and associative cortical inputs, potentially enabling enhanced sensorimotor and cognitive control ^11,65^.

**Figure 6.**
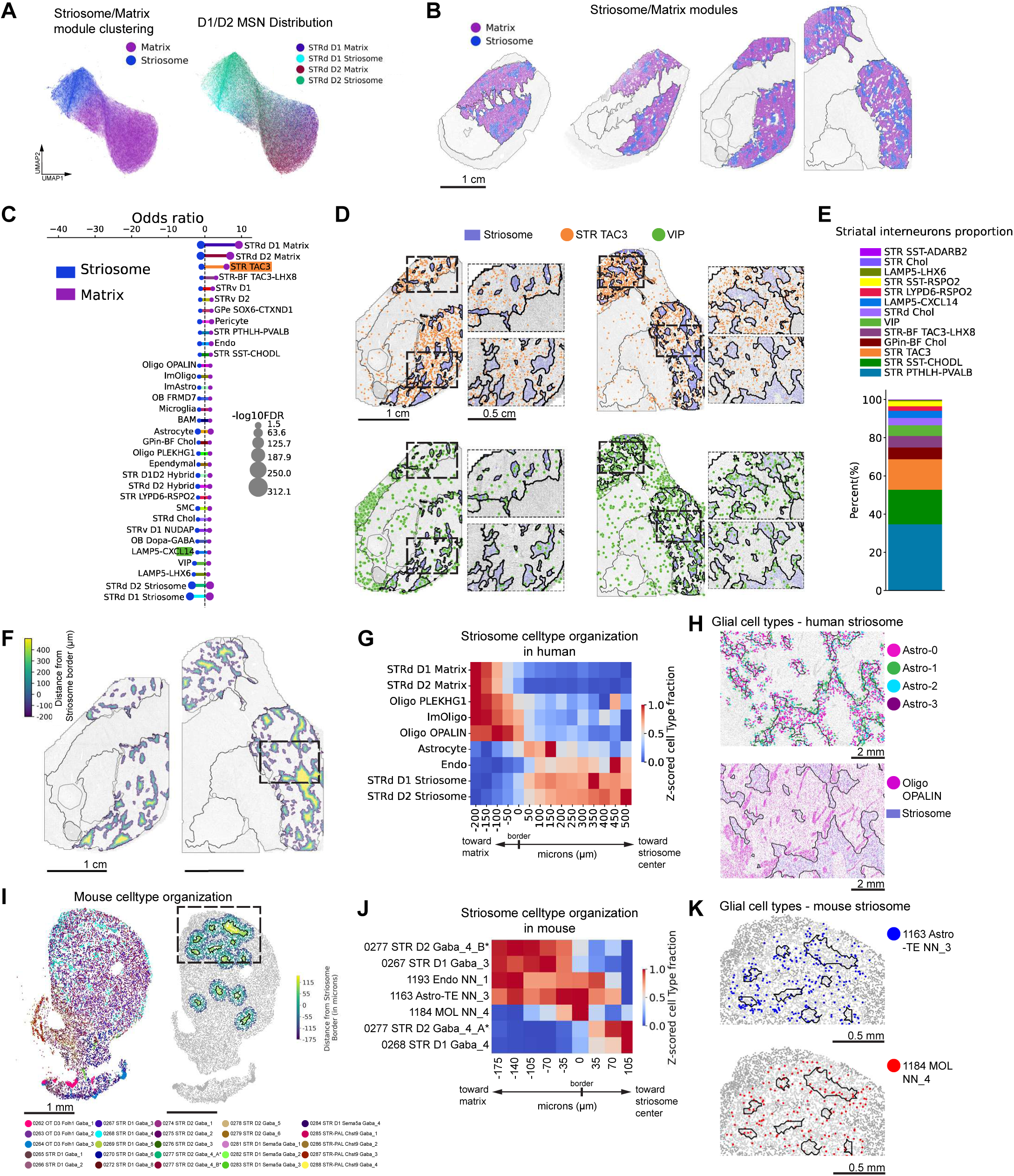
Cellular community and proximity analyses resolve striosome–matrix organization, gene-expression spatial gradients, and cross-species correspondence in the human basal ganglia. (**A**) UMAP computed based on cellular community analysis restricted to MSN cell types. Leiden clustering defines the matrix (purple) and striosome (blue) (left). Different types of MSNs are colored as indicated (right). (**B**) Spatial maps of striosome and matrix compartments (defined in **A**) across anterior–posterior (A–P) sections (AP 18a, 19a, 26a, 34a). (**C**) Cell-type enrichment (odds ratios) for striosome versus matrix compartments. Dot size reflects statistical significance (−log₁₀ FDR). P-values are derived using right-tailed chi-square tests; FDR adjustment via the Benjamini–Hochberg method; odds ratios via Cochran–Mantel–Haenszel test. For purpose of visualization, odds ratio values associated with the striosome are multiplied by −1. (**D**) Spatial distributions of interneuron subtypes showing compartmental biases: STR TAC3 GABAergic interneurons enriched in matrix and VIP+ GABAergic interneurons enriched in striosome regions. In the main text, STR TAC3 GABAergic interneurons are referred to as STR TAC3-PLPP4 using the long name of the cell type; lilac-blue shading marks striosome mask defined in (**A**). (**E**) Relative abundance of medial ganglionic eminence (MGE)– and caudal ganglionic eminence (CGE)–derived interneuron subtypes in the striatum. (**F**) Spatial heatmap of distance for each cell to the nearest striosome boundary (positive values = inside the striosome; negative values = outside the striosome). (**G**) Cell-type distributions as a function of distance to striosome boundaries, illustrating cell-type organization within striosomes in human. (**H**) Spatial distribution of four astrocyte subclusters defined based on the genes differentially expressed across striosome borders (top); and Oligo-OPALIN oligodendrocytes enriched at striosome borders (bottom). Panels show magnified views of the boxed region in (**F**). (**I**) Spatial distributions of mouse striatal neuronal subtypes re-plotted using the Allen Brain Institute Whole Mouse Brain MERFISH dataset (left). Cellular community analysis identifies mouse striosome–matrix compartments with the spatial heatmap of per-cell distance to the nearest striosome boundary (positive values = inside the striosome; negative values = outside the striosome) (right). (**J**) Cell-type distributions as a function of distance to striosome boundaries in mouse, showing compartmental organization analogous to human. (**K**) Enlarged view of the boxed region in mouse dorsal striatum (**I**) showing distributions of the border-enriched subtype of mouse astrocytes (Allen dataset: supertype: 1163 Astro-TE NN_3) (top) and oligodendrocytes (Allen dataset: supertype: 1184 MOL NN_4) (bottom) across striosome–matrix boundaries.\

Conversely, VIP+ and LAMP5+ interneurons display moderate enrichment within striosomal territories (Figure 6C, 6D bottom), although these populations are not exclusive to that compartment. To further explore the organization of cell types across the matrix-striosome boundary, we quantified the distance of each cell to the nearest striosome boundary and analyzed cell-type–specific density profiles as a function of proximity (Figure 6F). This analysis reveals astrocyte and Oligo OPALIN oligodendrocyte enrichment near striosome borders that form boundary-aligned microdomains (Figure 6G, 6H bottom). The Oligo OPALIN subtype, previously identified along striatopallidal fiber tracts (Figure 3), appears to delineate the perimeters of striosomal territories and highlights glial contributions to compartmental architecture.

To assess whether gene expression varies systematically with spatial proximity to striosomes, we examined transcript gradients along the striosome-to-matrix axis. Several astrocyte-enriched genes exhibit distance-dependent expression patterns (Figure S7A). Clustering based on these genes resolves four astrocyte subpopulations that occupy distinct positional zones—core, boundary, and perimatrix (Figures 6H, top); this organization is consistent across different sections (Figure S7B-E).

We next investigated cross-species conservation of this organization using the Allen Institute’s whole-mouse-brain MERFISH dataset ^10^. Orthologous mouse MSNs were identified by correlating gene-expression signatures with human subtypes (Figure S8A, B). Mouse supertypes 0268 STR D1 GABA_4 and 0267 STR D1 GABA_3 correspond to human STRd D1 Striosome and STRd D1 Matrix MSNs, respectively (Figure S8C–D). Similarly, finer subclustering of the mouse 0277 STR D2 Gaba_4 supertype, designated as A* and B*, map to human STRd D2 Striosome MSN and STRd D2 Matrix MSN, respectively (Figure S8E–F).

Using these orthologous MSN pairs, we delineated striosome–matrix boundaries in the mouse striatum and performed analogous proximity analyses in which we computed the distance of cells from border of nearest striosome (Figure 6I) and quantified the distribution of cells as a function of distance. The results are highly concordant across species: specific glial populations, Astro-TE NN_3 astrocytes and MOL NN_4 oligodendrocytes, are enriched at striosome borders in mice (Figure 6J-K).

These findings demonstrate conserved organizational principles and orthologous cellular microenvironments within striosome–matrix systems of the human and murine basal ganglia. The identification of matrix-confined TAC3–PLPP4 interneurons and boundary-enriched glial subtypes further reveals an evolutionarily elaborated compartmental architecture in the human striatum that integrates neuronal and non-neuronal components across spatial scales.

### Conserved molecular gradients and cross-species correspondence revealed by Stereo-seq mapping

While cell-type composition defines spatial modules, these modules ultimately emerge from underlying transcriptional programs. To directly examine these patterns, we applied unsupervised clustering to spatially weighted whole-transcriptome profiles derived from Stereo-seq data, independent of prior cell-type annotation (see Methods). This spatial analysis recapitulates the mesoscale modules identified by MERFISH+ cellular community analysis and aligns closely with established anatomical territories of the human basal ganglia (Figure 7A, Figure S9A).

**Figure 7.**
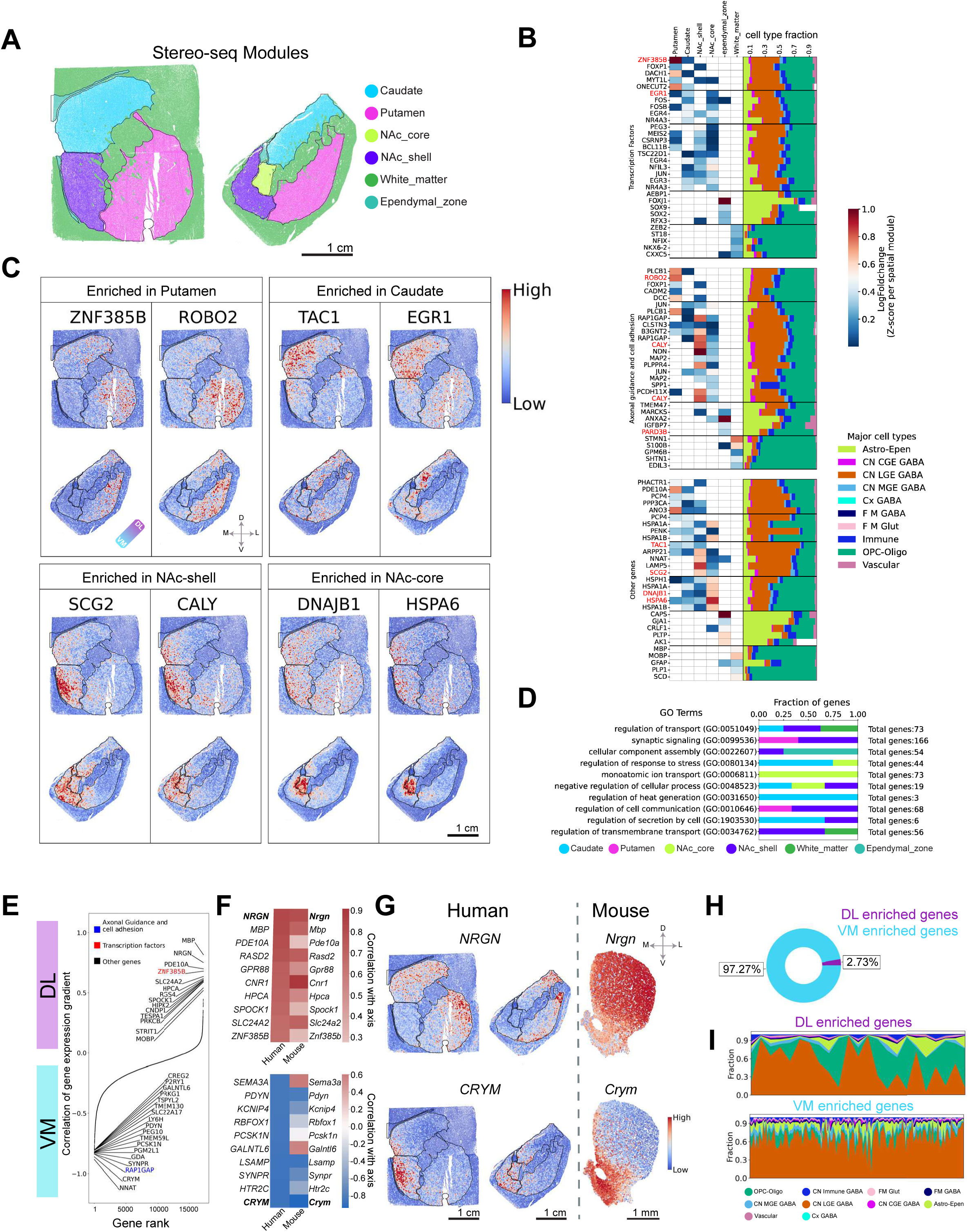
Stereo-seq spatial transcriptomics mapping of the human basal ganglia reveals conserved spatial gene-expression gradients between human and mouse. (**A**) Whole-transcriptome spatial modules mapped onto anatomical regions in sections AP 18a and AP 19a, the same color code as that depicted in Figure 5. (**B**) Heatmap of module-marker genes (rows) across spatial modules (columns), with corresponding cell-type composition shown alongside; values are cell type fraction, note red highlighted genes depicted in greater detail below. (**C**) Spatial expression maps of red-highlighted representative module-enriched genes in sections AP 18a and AP 19a. (**D**) Gene ontology (GO) enrichment analysis for each spatial module. The horizontal bar values show the fractions of genes associated with the color-coded regions for each term. (**E**) Correlation of gene expression gradient along the dorsolateral–ventromedial (DL–VM) axis. Genes are ranked based on their gradient expression. (**F**) Top 10 DL-expressed genes in human basal ganglia and their corresponding DL–VM gradient expression in mouse striatum (top). Top 10 VM-expressed genes in human basal ganglia and their corresponding VM-DL gradient expression in mouse striatum (bottom). These related correlations highlight the relatively conserved DL–VM gene-expression gradients across species. (**G**) Spatial gene expression maps for matching genes from panel **F** in human basal ganglia and mouse striatum. (**H**) Fractional distribution of DL-enriched versus VM-enriched genes in human basal ganglia. (**I**) Fraction plots of cell-type-specific expression for DL (top) and VM (bottom) enriched genes. See cell type color keys below.

Each spatial module is characterized by distinct sets of regionally enriched genes (Figure 7B-C, Figure S9B), indicating the molecular specialization among the subregions of the basal ganglia. For example, the putamen exhibits strong enrichment of ZNF385B (a zinc-finger transcription factor) and ROBO2 (an axonal guidance molecule), whereas the caudate nucleus shows elevated expression of TAC1(a neuropeptide precursor) and EGR1(an immediate early transcription factor). The nucleus accumbens shell is marked by vesicle associated molecules SCG2 and CALY, while the NAc core is enriched for stress-response genes HSPA6 and DNAJB1 (Figure 7C).

We further examined the transcriptional difference between the regions by performing gene ontology (GO) enrichment analysis on the differentially expressed genes. Pathways such as synaptic signaling and regulation of cell communication are mainly associated with putamen and NAc shell, while processes such as regulation of secretion by cell and regulation of response to stress prominently involve the caudate with some contribution from the NAc shell and NAc core respectively. Some GO terms are uniformly distributed among multiple regions; regulation of transport involves the caudate, NAc-shell, and white matter, and negative regulation of cellular process is linked to the caudate, NAc core and NAc shell (Figure 7D).

In our Stereo-seq data, the genes displaying regional specificity constitute only ∼5%; majority of the remaining genes are expressed throughout the striatum. This led us to ask whether gene expression gradients in the human striatum are similar to those known in mice and non-human primates^4,5,66^. To investigate topographical continuity of gene expression across species we focused on dorsolateral-ventromedial (DL-VM) expression gradients, an established organizational axis in the mouse striatum^14,66^. We find numerous genes in the human striatum that display graded expression along this axis (Figure 7E). For example, PDE10A, NRGN, and SLIT1 are dorsolaterally enriched, whereas NNAT, CRYM, and RAP1GAP exhibit ventromedial bias. Importantly, the majority of gradient-aligned genes show directionality concordance across species as illustrated by the similarity of the gradient scores (Figure 7F) and spatial expression maps (Figure 7G and Figure S9C) of genes between mouse and human. This underscores evolutionarily conserved transcriptional patterning that likely supports shared basal ganglia circuit architecture and function. In addition, we observe in both mouse and human, the expression gradient is dominated by the ventromedial direction (Mouse: 73%, Human: 97.2%) as opposed to the dorsolateral direction (Mouse: 26%, Human: 2.73%) (Figure 7H and Figure S9D).

To identify cellular contributors to these spatial gradients, we mapped gradient-associated genes onto cell types defined in the Allen Institute Basal Ganglia Consensus Cell Type Atlas. Most gradient-associated transcripts are enriched in lateral ganglionic eminence (CN/LGE)-derived GABAergic neurons, the principal MSN population. A subset of dorsolateral-enriched genes, however, are expressed within oligodendrocyte lineage (OPC–Oligo) cells, that reflects the dense myelinated fiber tracts that occupy dorsal striatal territories (Figure 7I). In contrast, in mouse we do not observe this; LGE neurons dominate in both directions (Figure S9E).

These analyses demonstrate that Stereo-seq captures transcriptome-wide spatial gradients that mirror functional and anatomical axes of the basal ganglia. The high concordance of these gradients between humans and mice highlights conserved molecular topographies that organize the macroscale regions across species (Figure 7F-I).

## DISCUSSION

By integrating the two high-throughput spatial transcriptomics platforms (MERFISH+ and Stereo-seq), we generated a multiscale spatial transcriptomic atlas of the human basal ganglia spanning 6 orders of magnitude in scale from subcellular transcript organization (∼50 nm) within single-cells to the multi-centimeter (∼4cm) spatial arrangement of cell types across basal ganglia nuclei. This large-scale atlas, which covers ∼7 million cells in total, encompasses all major nuclei including the caudate, putamen, nucleus accumbens, and the globus pallidus across four neurotypical postmortem brains. In addition to providing a comprehensive molecular and anatomical reference for the neuroscience community, our work establishes a scalable experimental and computational framework for performing large-area, high-resolution spatial transcriptomics in human tissue.

While much of the spatial organization of cell types and circuit domains in the human basal ganglia has been derived from rodent and non-human primate studies ^13,67^, our dataset provides the first comprehensive spatial and molecular reference directly in humans. The resulting framework reveals how subcellular gene expression, cell type identities and spatial cellular community structures may converge to shape human basal ganglia organization and potentially underlie its function.

### A unified molecular and spatial taxonomy across human basal ganglia

MERFISH+ resolves the spatial organization of 60 transcriptionally distinct cell types that includes the subclassification of eight major neuronal classes spanning anterior–posterior strata. Within the striatum, we map both the most abundant D1 and D2 MSNs into striosome and matrix compartments as well as the less abundant, hybrid-identity MSN cell types (e.g., STR D1D2 Hybrid; STRd D2 Hybrid), and show their preferential localization to the caudate, putamen, and NAc but not to GP or internal capsule. Within the globus pallidus, we delineate six GP-associated classes: GPi Shell, GPi Core, GPin-BF Cholinergic, CN LHX6–GBX1, GPe SOX6–CTXND1, and GPe MEIS2–SOX6 based on their single-cell transcriptional profiles and reveal their spatial locations within the GPe/GPi shell and core anatomical regions. We also find that non-neuronal classes are molecularly and spatially organized: the OPC–Oligo lineage comprises ∼60–70% of non-neuronal cells and partitions into COP, OPC, Oligo PLEKHG1, ImOligo, and Oligo OPALIN subtypes with distinct regional enrichments (Oligo PLEKHG1 in internal capsule/GP; Oligo OPALIN broadly distribute with enhanced abundance in striatopallidal fibers). Six astrocyte subtypes, distinguished by markers such as AQP4, AQP1, SLC6A11, CHI3L1, and ADGRV1, show region-specific topographies (e.g., WM_Astro/AQP1 in white matter; STR_Astro-2/ ADGRV1 in striatum; GP-TH_Astro/SLC6A11 in GP).

### Mesoscale cellular communities and striosome–matrix architecture

Our cellular community analysis based on the proximity of the 60 identified cell types reveals 14 spatial modules in the human basal ganglia sections, providing an unbiased, refined anatomical delineation of the imaged regions that aligns closely with the Allen Human Brain Atlas parcellation ^59^. The differential enrichment of cell-types across spatial modules recapitulates expected spatial segregations based on the Mammalian Basal Ganglia Consensus Cell Type Atlas reference (Allen Institute; RRID:SCR_024672) (e.g., GPe SOX6–CTXND1/MEIS2–SOX6 in GPe; GPi Core/Shell in GPi; STRv D1/D2 in NAc; STRd D1/D2 in the caudate/putamen) including regional interneuron biases: STR FS PTHLH–PVALB enriched in the caudate, ChAT+ enriched in the putamen, and FM GATA3–TCF7L2 concentrated in GPi. Within NAc, we uncover that both D1- and D2-striosomal MSNs populate NAc shell but are largely absent in core, consistent with TH (striosome) and KIRREL3 (matrix) compartment markers.

The matrix and striosome spatial modules derived from MSN subtype composition reveals fine-scale spatial organization biases. Specifically, TAC3–PLPP4 interneurons, which are expanded in primates ^64^, are matrix-restricted, whereas VIP+ and LAMP5+ interneurons are striosome-enriched but not exclusive. Across the striosome boundaries, we find enrichment of specific subpopulations of astrocytes and oligodendrocytes which delineates boundary-aligned glial microdomains. Cross-species comparison with the MERFISH-derived mouse cell type atlas ^10,14^ identifies orthologous MSN counterparts (e.g., mouse 0268 D1 and 0267 D1 supertypes corresponding to human STRd D1 striosome and matrix respectively) and confirms border-enriched astrocyte (Astro-TE NN_3) and oligodendrocyte (MOL NN_4) supertypes in mouse. These data argue for conserved compartment organization principles and orthologous boundary microenvironments between human and rodent striata, while also highlighting primate and human specializations (e.g., matrix-restricted TAC3–PLPP4) ^67,68^.

### Subcellular transcript localization informs morphology and projection architecture

MERFISH+ subcellular transcript localizations enable characterization of cell morphology and projection-inferred connectivity. Our 3D analysis of RNA transcript localization provides soma-volume measurements that span ∼5-fold variation across neuronal types. Although this strategy differs from traditional methods that offer more detailed cell morphology, including immunofluorescence staining and Cajal–Golgi impregnation methods ^69,70^, the key advantage of MERFISH+ based quantification is its ability to distinguish and measure dozens of neuronal subclasses at tissue-wide scale. We find that cholinergic interneurons and pallidal neuron subtypes (e.g., GPi Core/Shell; GPe NDB–SI–LHX6/8–GBX1; GPe MEIS2–SOX6) exhibit the largest soma volumes, whereas LAMP5+ interneurons and MSNs are among the smallest. These findings are supported by previous histological observations in basal ganglia ^51–53^. While the nine MSN subtypes show largely similar volumes, SST+ interneuron subclasses diverge markedly; for example, SST–CHODL versus SST–ADARB2 interneurons show substantial differences. These physical parameters could inform future neural network models in classifying cell types based on label free methods reporting on size and morphology ^71,72^.

Quantification of putative “neuritic” transcripts within the GPi and GPe recapitulates in human the canonical projection pathways measured directly in animal models. Specifically, D1-enriched genes (e.g., RELN, EBF1, SLIT1) are found to preferentially accumulate in GPi projections, whereas D2-enriched genes are biased to GPe projections. These correlation patterns recapitulate the direct (D1→GPi) versus indirect (D2→GPe) pathway topology in rodent and non-human primate models ^3,6,68^. Thus, subcellular transcript “geography” provides a quantitative readout of both cell size/shape and projection-inferred circuitry in human brain tissue.

### Genome-wide transcriptional modules and a conserved DL-VM gradient

Unsupervised clustering of the whole-transcriptome spatial distribution (based on Stereo-seq) recapitulates MERFISH+ cell-type community modules and classical anatomy and provides novel gene enrichment within each anatomical subregions. We find that a dorsolateral- ventromedial gradient emerges as a dominant axis of transcriptional variation at the multi-centimeter-scale; this axis is conserved in the mouse at the multi-millimeter scale ^10,14^ revealing dorsolateral-biased genes such as PDE10A, NRGN and SLIT1 and ventromedial-biased genes such as NNAT, CRYM, RAP1GAP. Most of the genes exhibiting the gradient are expressed in CN LGE GABA neurons, with a dorsal contribution from OPC–Oligo lineages, consistent with differential myelination in the dorsal fiber bundles ^73,74^.

### Functional implications

Functionally, the underlying cell types, cellular communities, subcellular transcript geography, and conserved spatial gradients suggest an intricate multi-layered organizational architecture for neurotypical human basal ganglia ^75,76^. The finding that subcellular RNA localization captures projection-inferred signatures suggests a promising avenue for inferring human circuit topology directly from spatial transcriptomic data - an approach that circumvents the limitations of traditional neural tracing in human tissue. In addition, astrocyte and oligodendrocyte niches aligned with striosome boundaries point to specialized metabolic and myelination interfaces at these compartmental perimeters. Finally, the primate-expanded, matrix-restricted TAC3–PLPP4 interneurons may reflect evolutionary adaptations to the enlarged prefrontal cortex, potentially supporting the increased computational demands of human-specific action selection, valuation, and cognitive flexibility ^11^.

Clinically, our dataset highlights specific neuronal and glial subtypes including GPi core versus shell populations and boundary-enriched glial communities, as potential substrates of selective vulnerability in movement and neuropsychiatric disorders. Establishing the neurotypical abundance and spatial distribution of D1, D2, and D1/D2 hybrid MSN subtypes in the caudate, putamen, and nucleus accumbens provides a crucial baseline for detecting disease-associated shifts in cell number, composition, and transcriptional state. Such comparisons will be particularly valuable for elucidating cell-type-selective vulnerability in basal ganglia disorders, including Huntington’s disease and Parkinson’s disease ^77,78^.

### Limitations of the study

This study is limited by the number and demographic diversity of available donors, as well as inherent constraints associated with postmortem human tissue. Although our multimodal spatial framework provides a comprehensive baseline, additional orthogonal validations such as immunohistochemistry, in situ sequencing, and in vivo electrophysiological or connectivity assays would further strengthen cell-type and spatial community assignments. Future work should extend this approach to a broader range of ages and demographic backgrounds and apply this methodology to comparative analyses of neurological and psychiatric disorders in human tissue.

In summary, by coupling genome-wide spatial profiling with subcellular transcript localization and community-level modeling, we deliver a multiscale, molecular atlas of human basal ganglia organization from single transcripts to cellular communities and anatomical modules that shows features unique to humans and those that are conserved across species. This atlas establishes a foundational framework for studying human basal ganglia architecture and its disruption in neuropsychiatric and neurodegenerative disorders, enabling cross-species and disease-state comparisons at single-cell and community scales.

## ACKNOWLEDGEMENTS

We thank members of the Xu laboratory for their valuable feedback and discussion throughout this project. We are grateful to Dr. Ruth Walker for her insightful comments. We also acknowledge Drs. Elizabeth and Edwin Monuki, along with our UCI colleagues, for their contributions to establishing the UCI BICAN Brain Bank and supporting the collection of neurotypical human brain samples. This work was in part funded by the NIH grant [UM1MH130994 Center for Multiomic Human Brain Cell Atlas] and UC Irvine Center for Neural Circuit technology development funds. This publication was supported and coordinated through the Brain Initiative Cell Atlas Network (BICAN, RRID:SCR_022794).

## AUTHOR CONTRIBUTIONS

X.X. and B.B. conceived the study. X.X., B.B. and Z.T designed the experiments. B.T.B., B.B., and X.X. designed the gene probe libraries. J.B., P.A.S.M, G.W, and Z.T. prepared samples, Z.T. and S.C.D performed the experiments. B.T.B., B.E., R.L., and B.B. performed data analysis. X.X., and B.B. interpreted data. B.T.B., Z.T., T.C.H., B.B. and X.X. wrote and edited the manuscript with input from G.W., S.C.D., Q.Y., R.L., M.N., S.P., Q.Z., M.B., and J.E. X.X. supervised the study.

## Supplementary Materials

**Supplementary Movie 1. MERFISH+ spatial transcriptomics of a large human basal ganglia section visualized across scales.** The movie depicts successively greater magnified views starting from multi-centimeter tissue architecture to subcellular resolution scale, revealing transcriptionally distinct cell types and local cellular neighborhoods, and individual RNA transcripts visualized at submicrometer precision (color-coded by gene identity).

**Table S1. Human brain sample information.** Detailed information on human donor identification number, age, sex, race/ethnicity, postmortem interval (PMI) from time of death to tissue freezing, RNA integrity number (RIN) indicating sample quality, cause of death, section AP, subregions profiled.

**Supplemental Data** for the information related to the MERFISH+ encoding probes, adaptor probes, and readout probes and the gene panel codebook.

## Supplemental Figure Legends

**Figure S1. Comparative spatial maps of major cell types of adjacent coronal sections of human basal ganglia between MERFISH+ and Stereo-seq. (A)** Comparative spatial distributions of Astro-Epen, CN CGE GABA, CN LGE GABA, CN MGE GABA, Immune, and OPC-Oligo cell types mapped by MERFISH+ and Stereo-seq, respectively, showing strong agreement between two approaches. **(B)** Pearson correlations of per-cell count or mean gene expression between MERFISH+ and Stereo-seq show significant correlation between measurements, see panels for correlation values. **(C)** Pearson correlations of per-cell count or mean gene expression between independent MERFISH+ assays (MER1 vs. MER2/3/4), demonstrate significant reproducibility across platforms and replicates. MER1/2/3/4 = MERFISH+ assays 1/2/3/4, see panels for correlation values. **(D)** Pearson correlations of mean gene expression between independent Stereo-seq experiments show significant correlations. Astro-Epen = astrocyte–ependymal; CN CGE GABA = cerebral nuclei caudal ganglionic eminence GABA; CN LGE GABA = cerebral nuclei lateral ganglionic eminence GABA; CN MGE GABA = cerebral nuclei medial ganglionic eminence GABA, see panels for correlation values. Scale bar = 1 cm.

**Figure S2. Graphical representation of top differentially expressed genes between the 60 transcriptomics cell types identified by MERFISH+.** The dot-plot shows 150 differential expressed genes out of the 673 MERFISH+ imaged genes, thus validating the distinction between 60 transcriptomics cell types.

**Figure S3. Spatial distributions and relative abundances of non-MSN and globus pallidus–associated neuronal subtypes across anterior–posterior coronal sections of the human basal ganglia.** (A) Relative proportions of all neuronal cell types including non-MSN and globus pallidus (GP) neuron subtypes—as well as CGE-and MGE-derived GABAergic neuron subclasses, in both combined and individual MERFISH+ assays, showing all sections versus AP-specific proportions. (B) Spatial distributions of D1+ and D2+ medium spiny neurons (MSNs) across anterior–posterior sections of the human basal ganglia. (C) Spatial distributions of diverse non-MSN neuronal subtypes, including VIP GABA, LAMP5–CXCL14 GABA, STR SST–ADARB2 GABA, STR SST–RSPO2 GABA, STR SST–CHODL GABA, STR LYPD6–RSPO2 GABA, STRd Cholinergic GABA, OT D1ICj, LAMP5–LHX6 GABA, STR TAC3–PLPP4 GABA, OB FRMD7 GABA, and STR–BF TAC3–PLPP4–LHX8 GABA cell types show subregional expression specificity. Scale bar = 1 cm.

**Figure S4. Spatial distributions of GPi/GPe neuron subtypes and associated marker expression in the human globus pallidus.** (**A**) Relative proportions of forebrain–midbrain (FM) GABAergic and FM glutamatergic neuron subtypes identified in MERFISH+ data, including all sections combined and specific AP sections. (**B**) Spatial distributions of AMY–SLEA–BNST GABA, GPe MEIS2–SOX6 GABA, AMY–SLEA–BNST D1 GABA, and GPin–BF Cholinergic GABA neuron subtypes across globus pallidus. (**C**) Heatmap showing expression of top marker genes distinguishing six GPi/GPe neuron subtypes. (**D**) Spatial maps showing representative marker genes for each of the six GPi/GPe neuron subtypes.

**Figure S5. Spatial distributions of non-neuronal cell sub-types in the human basal ganglia.** (**A**) Relative proportions of astrocyte–ependymal (Astro-Epen) subtypes, vascular-associated subtypes, and immune subtypes across all MERFISH+ assays, including all sections combined and AP sections 18-34, note AP19 includes the greatest ependymal zone surface abutting the ventricle. (**B**) Spatial maps of non-neuronal subtypes, including Oligo PLEKHG1, oligodendrocyte precursor cells (OPCs), Oligo OPALIN, committed oligodendrocyte precursors (COPs), immune-associated oligodendrocytes (ImOligo), astrocytes, immune-associated astrocytes (ImAstro), ependymal cells, vascular leptomeningeal cells (VLMCs), smooth muscle cells (SMCs), pericytes, endothelial cells, microglia, monocytes, B cells, and T cells from AP18-34. Scale bar = 1 cm for all panels.

**Figure S6. Spatial maps of oligodendrocyte and astrocyte sub-types in the human basal ganglia.** (**A**) Spatial distributions of two oligodendrocyte subtypes (Oligo OPALIN and Oligo PLEKHG1) in anterior and posterior sections of the human basal ganglia. (**B**) Spatial subregion specific distributions of six astrocyte subtypes across anterior–posterior sample sections. (**C**) Spatial expression maps of representative marker genes that correspond to each astrocyte subtype across anterior–posterior sample sections show subregion specific expression.

**Figure S7. Astrocyte subclusters stratified by gene-expression gradients across striosome–matrix boundaries.** (**A**) Heatmap of astrocytic gene expression as a function of distance from the striosome–matrix border. Distance is measured relative to the border (center), where 0 represents the striosome-matrix border, and spans −500 µm into matrix regions (negative values) and +500 µm into striosome interiors (positive values). (**B**) UMAP embedding computed from clustering of astrocytes located within ±500 µm from the striosome-matrix border based on genes shown in (**A**) in the posterior sample AP26a (left) and the spatial map showing the distribution of the subclusters relative to striosome boundaries (middle). Right: Zoomed-in view of the boxed region (left). (**C**) Same analysis as in (B), shown for posterior section AP34a: UMAP embedding (left), spatial distribution along striosome boundaries (middle), and zoom-in of the boxed region (right). **(D)** Density distribution of astrocyte subclusters as a function of distance from the striosome–matrix border in AP26a, 0 represents the striosome-matrix border. **(E)** Density distribution of astrocyte subclusters as a function of distance from the striosome–matrix border in AP34a, 0 represents the striosome-matrix border.

**Figure S8. Cross-species mapping between human and mouse orthologous basal ganglia cell types.** (**A**) Corresponding section position in the Allen Institute for Brain Science (AIBS) CCF v3 mouse reference atlas (left), and the spatial transcriptomics section from the AIBS Whole Mouse Brain (WMB) MERFISH C57BL/6J-638850 dataset with annotated regional parcellation (right). The red box indicates the region used to extract basal ganglia–associated cell types. (**B**) Cell-type expression correlation matrix comparing human and mouse basal ganglia cell types. DRD-related cell types are highlighted (red text and bounding box). Columns = mouse cell types; rows = human cell types. 0277 STR D2 Gaba_4_A* and 0277 STR D2 Gaba_4_B* represent two manually combined mouse cell types derived from higher-resolution clusters of 0277 STR D2 Gaba_4. (**C–F**) Fine-grained mapping of human STRd D1 striosome MSNs (**C**), D1 matrix MSNs (**D**), D2 striosome MSNs (**E**), and D2 matrix MSNs (**F**) to orthologous mouse cell types. Top: gene-expression correlation and normalized marker-gene expression profiles. Rows = human and mouse cell types (mouse ordered by correlation to the human type); columns = marker genes from the human subtype sorted by relative abundance. Mouse expression reflects corresponding orthologous gene values. Middle: spatial distributions of the human MSN cell types. Bottom: spatial distributions of the best-matched orthologous mouse MSN cell types.

**Figure S9. More spatial module analysis using additional human basal ganglia Stereo-seq data. (A)** Whole-transcriptome spatial modules identified in additional Stereo-seq samples including the external segment of the globus pallidus (GPe). **(B)** Spatial expression maps of representative region-enriched genes: PDE10A (putamen), PCP4 (caudate), PEG10 (NAc), and SLC6A11 and PVALB (GPe). **(C)** Spatial expression map showing conserved dorsolateral–ventromedial (DL–VM) gradients in human and mouse striatum. DL-enriched genes: CNR1 and SRRIT1; VM-enriched genes: SYNPR and PDYN. **(D)** Fractional distribution of DL-enriched versus VM-enriched genes in mouse striatum. **(E)** Fraction plots of cell-type-specific expression for DL (top) and VM (bottom) direction in mouse striatum.

## STAR* METHODS

### KEY RESOURCES TABLE

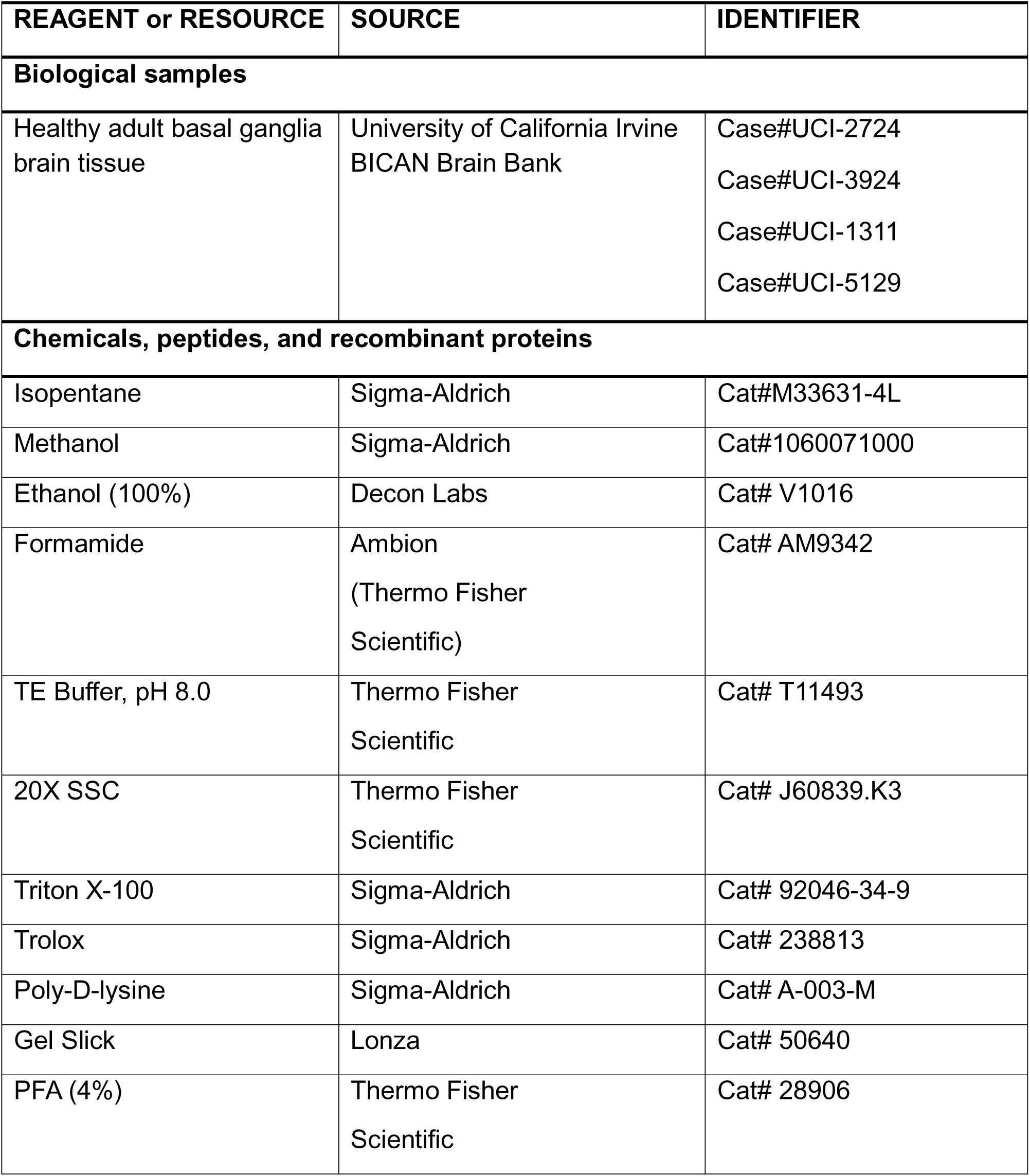
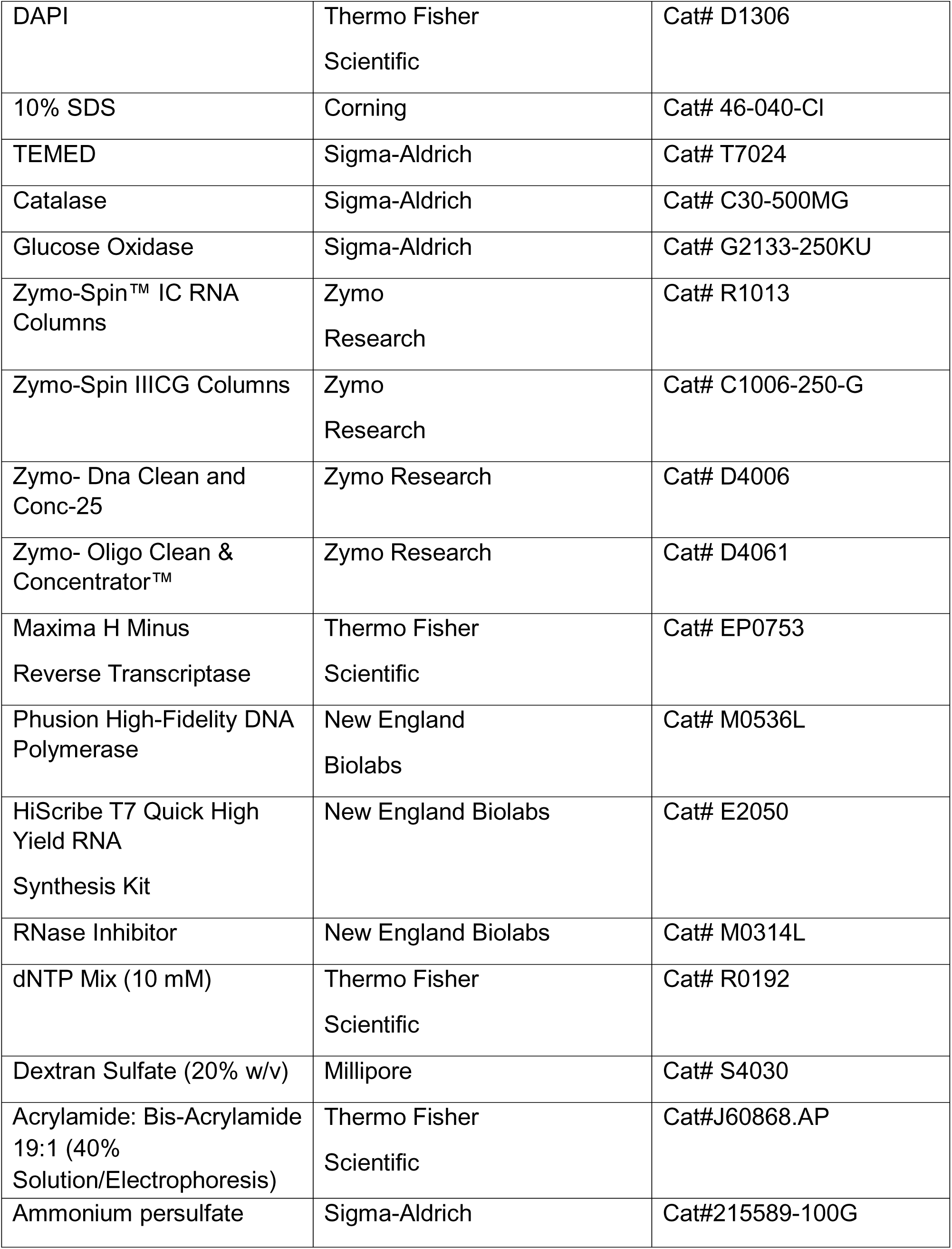
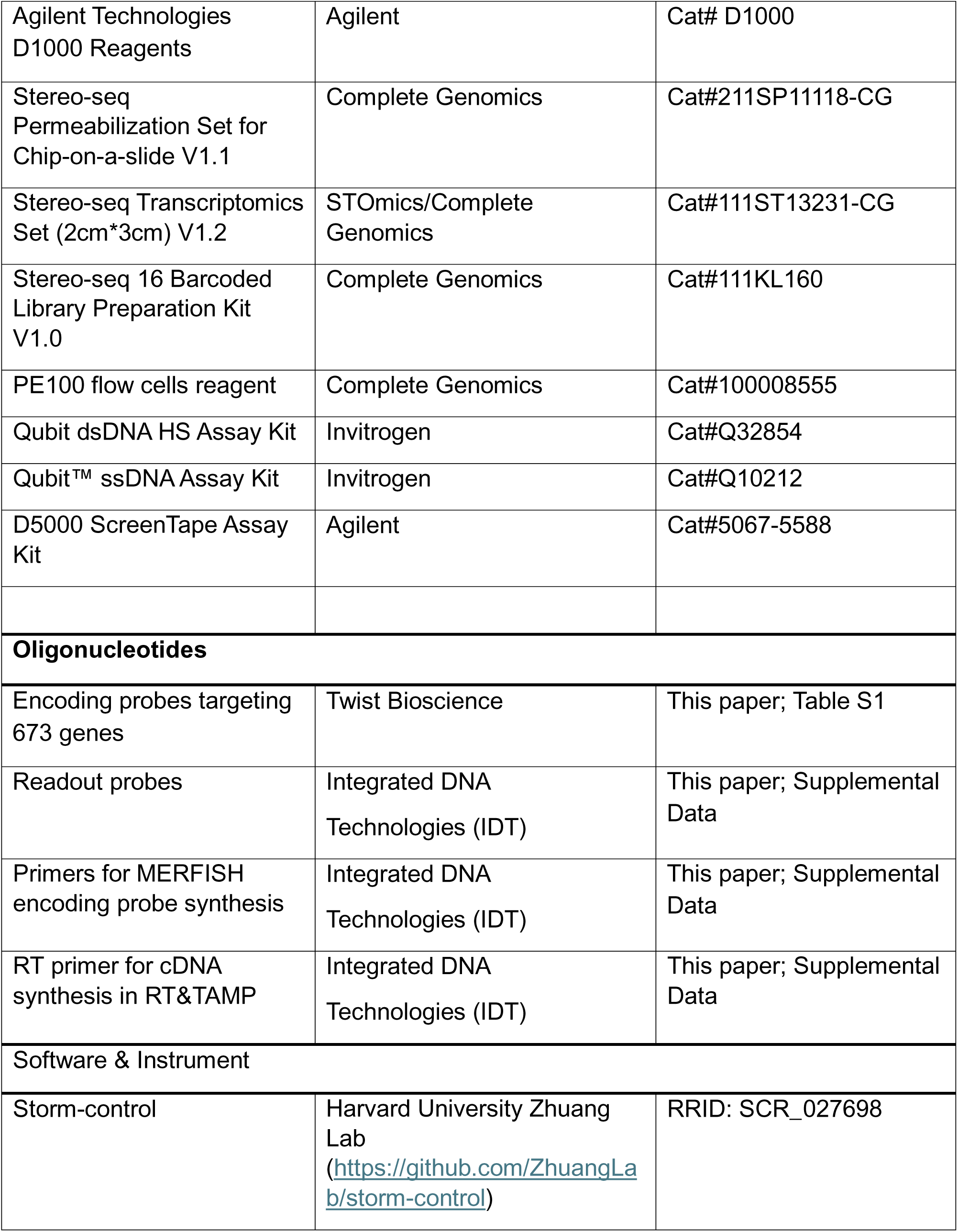
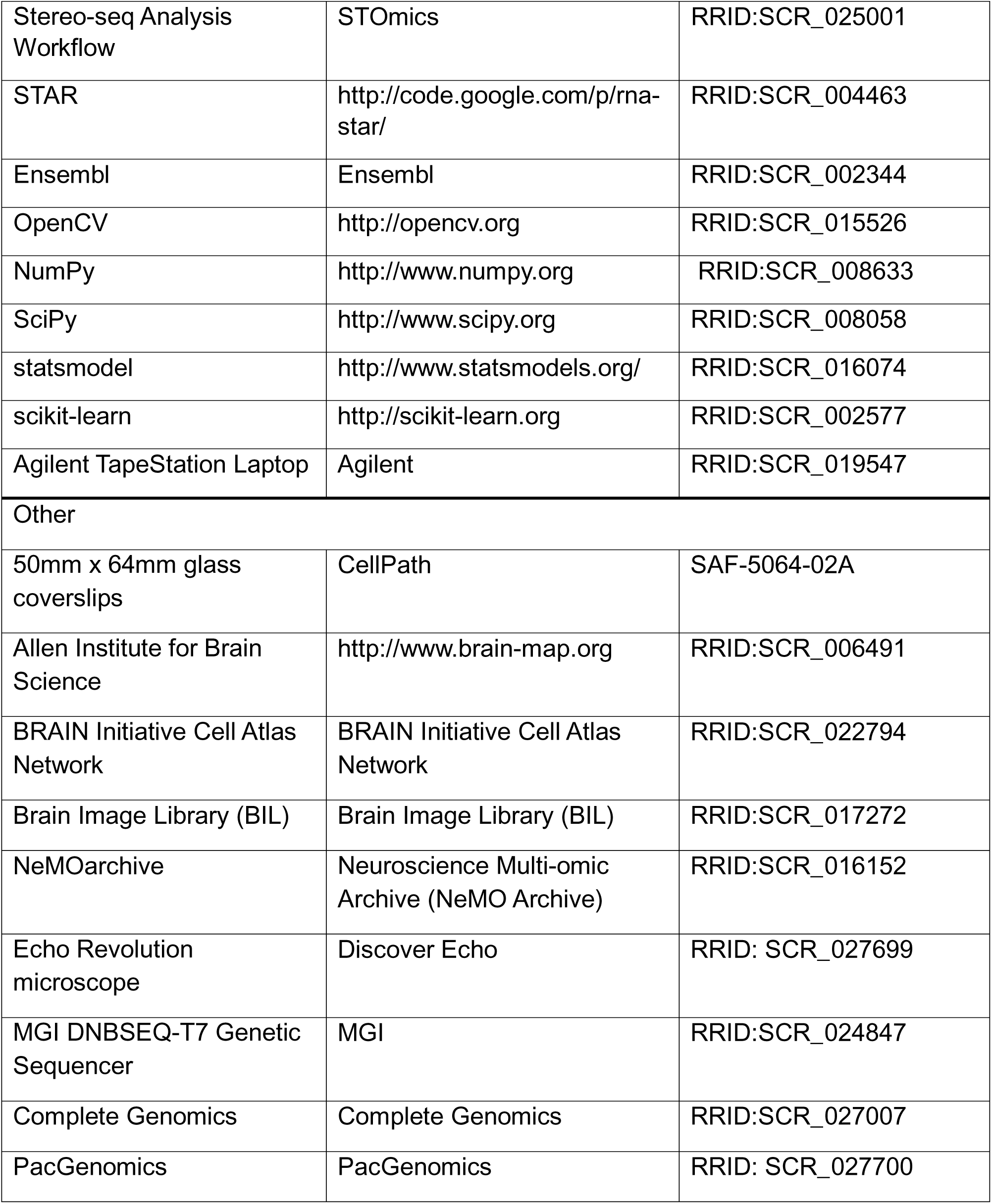

| REAGENT or RESOURCE | SOURCE | IDENTIFIER |
| --- | --- | --- |
| **Biological samples** |  |  |
| Healthy adult basal ganglia brain tissue | University of California Irvine<br>BICAN Brain Bank | Case#UCI-2724<br>Case#UCI-3924<br>Case#UCI-1311<br>Case#UCI-5129 |
| **Chemicals, peptides, and recombinant proteins** |  |  |
| Isopentane | Sigma-Aldrich | Cat#M33631-4L |
| Methanol | Sigma-Aldrich | Cat#1060071000 |
| Ethanol (100%) | Decon Labs | Cat# V1016 |
| Formamide | Ambion<br>(Thermo Fisher Scientific) | Cat# AM9342 |
| TE Buffer, pH 8.0 | Thermo Fisher Scientific | Cat# T11493 |
| 20X SSC | Thermo Fisher Scientific | Cat# J60839.K3 |
| Triton X-100 | Sigma-Aldrich | Cat# 92046-34-9 |
| Trolox | Sigma-Aldrich | Cat# 238813 |
| Poly-D-lysine | Sigma-Aldrich | Cat# A-003-M |
| Gel Slick | Lonza | Cat# 50640 |
| PFA (4%) | Thermo Fisher Scientific | Cat# 28906 |
| DAPI | Thermo Fisher Scientific | Cat# D1306 |
| 10% SDS | Corning | Cat# 46-040-CI |
| TEMED | Sigma-Aldrich | Cat# T7024 |
| Catalase | Sigma-Aldrich | Cat# C30-500MG |
| Glucose Oxidase | Sigma-Aldrich | Cat# G2133-250KU |
| Zymo-Spin™ IC RNA Columns | Zymo Research | Cat# R1013 |
| Zymo-Spin IIICG Columns | Zymo Research | Cat# C1006-250-G |
| Zymo- Dna Clean and Conc-25 | Zymo Research | Cat# D4006 |
| Zymo- Oligo Clean & Concentrator™ | Zymo Research | Cat# D4061 |
| Maxima H Minus Reverse Transcriptase | Thermo Fisher Scientific | Cat# EP0753 |
| Phusion High-Fidelity DNA Polymerase | New England Biolabs | Cat# M0536L |
| HiScribe T7 Quick High Yield RNA Synthesis Kit | New England Biolabs | Cat# E2050 |
| RNase Inhibitor | New England Biolabs | Cat# M0314L |
| dNTP Mix (10 mM) | Thermo Fisher Scientific | Cat# R0192 |
| Dextran Sulfate (20% w/v) | Millipore | Cat# S4030 |
| Acrylamide: Bis-Acrylamide 19:1 (40% Solution/Electrophoresis) | Thermo Fisher Scientific | Cat#J60868.AP |
| Ammonium persulfate | Sigma-Aldrich | Cat#215589-100G |
| Agilent Technologies D1000 Reagents | Agilent | Cat# D1000 |
| Stereo-seq Permeabilization Set for Chip-on-a-slide V1.1 | Complete Genomics | Cat#211SP11118-CG |
| Stereo-seq Transcriptomics Set (2cm*3cm) V1.2 | STOmics/Complete Genomics | Cat#111ST13231-CG |
| Stereo-seq 16 Barcoded Library Preparation Kit V1.0 | Complete Genomics | Cat#111KL160 |
| PE100 flow cells reagent | Complete Genomics | Cat#100008555 |
| Qubit dsDNA HS Assay Kit | Invitrogen | Cat#Q32854 |
| Qubit™ ssDNA Assay Kit | Invitrogen | Cat#Q10212 |
| D5000 ScreenTape Assay Kit | Agilent | Cat#5067-5588 |
| **Oligonucleotides** |  |  |
| Encoding probes targeting 673 genes | Twist Bioscience | This paper; Table S1 |
| Readout probes | Integrated DNA Technologies (IDT) | This paper; Supplemental Data |
| Primers for MERFISH encoding probe synthesis | Integrated DNA Technologies (IDT) | This paper; Supplemental Data |
| RT primer for cDNA synthesis in RT&TAMP | Integrated DNA Technologies (IDT) | This paper; Supplemental Data |
| **Software & Instrument** |  |  |
| Storm-control | Harvard University Zhuang Lab<br>(https://github.com/ZhuangLab/storm-control) | RRID: SCR_027698 |
| Stereo-seq Analysis Workflow | STOmics | RRID:SCR_025001 |
| STAR | http://code.google.com/p/rna-star/ | RRID:SCR_004463 |
| Ensembl | Ensembl | RRID:SCR_002344 |
| OpenCV | http://opencv.org | RRID:SCR_015526 |
| NumPy | http://www.numpy.org | RRID:SCR_008633 |
| SciPy | http://www.scipy.org | RRID:SCR_008058 |
| statsmodel | http://www.statsmodels.org/ | RRID:SCR_016074 |
| scikit-learn | http://scikit-learn.org | RRID:SCR_002577 |
| Agilent TapeStation Laptop | Agilent | RRID:SCR_019547 |
| Other |  |  |
| 50mm x 64mm glass coverslips | CellPath | SAF-5064-02A |
| Allen Institute for Brain Science | http://www.brain-map.org | RRID:SCR_006491 |
| BRAIN Initiative Cell Atlas Network | BRAIN Initiative Cell Atlas Network | RRID:SCR_022794 |
| Brain Image Library (BIL) | Brain Image Library (BIL) | RRID:SCR_017272 |
| NeMOarchive | Neuroscience Multi-omic Archive (NeMO Archive) | RRID:SCR_016152 |
| Echo Revolution microscope | Discover Echo | RRID: SCR_027699 |
| MGI DNBSEQ-T7 Genetic Sequencer | MGI | RRID:SCR_024847 |
| Complete Genomics | Complete Genomics | RRID:SCR_027007 |
| PacGenomics | PacGenomics | RRID: SCR_027700 |

### RESOURCE AVAILABILITY

#### Material availability

Oligonucleotide probe sequences used for imaging can be found in Supplemental Data. These probes or materials for making these probes can be purchased from commercial sources, as detailed in the Key Resources Table.

#### Data and code availability

Image and sequencing data are in the process of being deposited at the Brain Image Library (BIL, RRID:SCR_017272) and the Neuroscience Multi-omic Archive (NeMO Archive, RRID:SCR_016152). The relevant data analysis codes are uploaded to Github under the repository: https://github.com/BBerackey/BICAN_human_basal_ganglia_spatial_transcriptomics_manuscript_2025

### EXPERIMENTAL DETAILS

#### Postmortem human basal ganglia samples

Four human brain cases with the basal ganglia were obtained from the University of California Irvine (UCI) BICAN brain bank under approval of the Institutional Review Board (IRB). These adult human postmortem brain samples were from white males of 36 to 53 years old and showed no evidence of neurodegenerative pathology (Table S1).

#### Preparation of multi-centimeter tissue sections for Stereo-seq and MERFISH+

To facilitate multimodal investigation of the same donor sample, each pre-dissected postmortem human coronal brain slab, which is 5–10 mm thick and containing the basal ganglia, was used in this study. To prepare adjacent brain sections for experiments, a tissue block measuring up to 4 cm × 5 cm and encompassing basal ganglia components was further dissected out of each coronal brain slab, quickly embedded in OCT embedding medium, and frozen in dry ice prechilled isopentane (SigmaAldrich, M33631). Cryosections of 10 µm and 16 µm thickness were then obtained using a Leica CM1850 cryostat at –18 to –20 °C and mounted onto Stereo-seq chips and specially coated 50 mm × 64 mm rectangular glass slides for MERFISH+, respectively. To fit the 2 cm × 3 cm Stereo-seq chips (STOmics/Complete Genomics, 111ST13231-CG), each large brain section was trimmed and/or subdivided within the cryostat to approximately 18 mm × 28 mm dimensions (Figure 1).

#### Sample processing for Stereo-seq

For Stereo-seq, tissue fixation and spatial transcriptomics procedures were conducted in accordance with the manufacturer’s protocol and established methods. Briefly, tissue sections mounted on Stereo-seq chips were dried at 37°C (5 min), then fixed in pre-cooled methanol (SigmaAldrich, 1060071000) for 30 mins at −20 °C. After fixation, sections were stained with Qubit™ ssDNA Assay Kit (Invitrogen, Q10212) and incubated in the dark for 5 mins, washed with 0.1× SSC buffer with 5% RNase inhibitor (0.05 U/mL, New England Biolabs, M0314L), the nuclei-stained tissue section was imaged using the FITC channel under a 10x microscope objective of Echo Revolution microscope (Discover Echo, RRID: SCR_027699). The resulting TIFF file was run through StereoMap ImageQC. After passing ImageQC, the chip was washed with 0.1× SSC before proceeding with permeabilization.

Permeabilization was conducted using the manufacturer’s buffer (Complete Genomics, 211KT13114) at 37°C for 12 minutes. RNA captured on Stereo-seq chips from permeabilized tissue was reverse transcribed at 42°C. The tissue was then removed, and cDNA was released from the chip using a transcriptomics reagent kit (Complete Genomics, 211KT13114). After reverse transcription, tissue removal and cDNA release and cDNA collection were done at 55°C for 3 to 18 hours. After size selection, amplification, and purification, cDNA concentrations were measured with the Qubit dsDNA HS Assay Kit (Invitrogen, Q32854). The distribution of cDNA fragment size was also measured using Agilent’s D5000 ScreenTape assay (Agilent, 5067-5588). For library construction, 20 ng of cDNA per sample was processed with a library preparation kit (Complete Genomics, 111KL160) followed by DNA nanoball (DNB) generation. Sequencing was performed on the DNBSEQ™ T7 platform at PacGenomics (Agoura Hill, CA, USA) with PE100 flow cells and 50 bp read 1 and 100 bp read 2 (Complete Genomics, 100008555).

### MERFISH+ design and methodology

#### 1. Gene selection

First, we compiled 992 candidate genes that comprised 150 canonical cell type marker genes, 20 control genes (housekeeping genes) and 822 genes that were differentially expressed between Group level cell types in the reference Human basal ganglia snRNA-seq dataset from the Allen Brain Institute (RRID:SCR_006491). We then refined the list by removing non-coding genes, and genes that are highly abundant in all cell types as they can lead to molecular crowding. We considered genes to be highly abundant, if their mean count across cell types (Group level) was greater than 20. After this step a total of 813 genes were remaining. Next, we identified the exon sequence from the longest isoform of each gene, and then we quantified the homology between the sequences based on a 12-nucleotide hash-table. Genes were removed if the fraction of homology was greater than 50% with p-value less than 0.001. Based on this criteria, 149 genes were removed, leaving 673 final candidate genes.

#### 2. Primary encoding probe, adaptor probe, and readout probe design

The MERFISH library was designed using a similar pipeline described in previous studies ^42–44^. As therein, each primary encoding probe contains a 40-mer target sequence that is complementary to the mRNA sequence of a gene, three 20nt readout sequences unique for each gene and two 20nt PCR primers. However, we made one modification to this design by adding a 4th readout sequence in order to help us to troubleshoot MERFISH experiments through smFISH imaging. To design the primary probes, we first generated overlapping 40-mer target sequences from the mRNA sequence of each gene. Next, we filtered the target sequences based on off-target binding score and other predefined screening requirements such as GC content, melting temperature, and homology against the genome, transcriptome and repetitive sequences. For each gene, we retained 10-150 40-mer target sequences that met the criteria, then we concatenated each sequence to 4 readout and 2 primer sequences. Finally, we screened the concatenated sequences for homology against the whole genome and excluded primary probes that had homology scores beyond 99 percentiles.

The selection of readout sequences that were concatenated to a target sequence of a gene was guided by a MERFISH codebook, which is a set of unique binary barcodes that determines in which imaging round the fluorescence signal of each gene will show up. We designed the codebook by first generating all possible 4 hamming distance binary words of length N = 28 using the set cover algorithm ^29^. Next, we used the Metropolis Hastings optimization algorithm to assign each binary word to a particular gene while minimizing the chances of molecular(signal) crowding due to shared channels when imaging genes that are co-expressed in the same cell type.

As described in the method section for MERFISH imaging, the primary probes were used in combination with adapter probes and readout probes during each imaging round. Adaptor probes are associated with a particular gene and they are made of 20nt sequence reverse complementary to the readout sequence used in the primary probe of the associated gene. In addition, they contain two identical sequences that serve as binding sites for readout probes. The readout probes are 20nt sequences that are conjugated with a fluorescence dye at the 5’ or 3’ end.

#### 3. Encoding probe synthesis

Primary encoding probes generation was performed from oligonucleotide pools as described previously ^42–44^. Oligonucleotide pools and primers were purchased from Twist Biosciences and Integrated DNA Technologies (IDT), respectively. Oligo pools were amplified through PCR with approximately 18 cycles using Phusion Plus PCR Master Mix (Thermo Fisher Scientific, F631L). The resulting PCR product was purified and concentrated with Zymo DNA clean & concentrator-25 (Zymo Research, D4006). RNA was generated from the purified PCR-amplified DNA through in-vitro transcription (IVT) using HiScribe T7 ARCA mRNA kit (NEB, E2060S).

Next, single strand DNA (ssDNA) was generated from the IVT-derived RNA through reverse transcription (RT) using Thermo Scientific Maxima Reverse Transcriptase (Thermo Fisher Scientific, EP0743). The RT product went through alkaline hydrolysis (1:1 of 1M NaOH and 0.5M EDTA) to remove RNA followed by ssDNA (encoding probes) purification using Zymo Oligo Clean & Concentrator (Zymo Research, D4061). The concentration and purity of the encoding probes were determined through NanoDrop while the size of the encoding probes was verified through Agilent TapeStation (RRID:SCR_019547). During the RT step, the forward primer introduced an acrydite anchor into the encoding probes which allowed the encoding probes to covalently bind with the polyacrylamide gel post encoding probes hybridization.

#### 4. Adaptor and readout probe preparation

Adaptor probes (total 28 adaptors) and readout probes (total 3 readout probes each tagged with one of the three fluorophores: 5Alex750N/3AlexF750N, 5Cy5/3Cy5Sp, 5Cy3/3Cy3Sp) were purchased from IDT. Adaptor probes and readout probes were diluted with a pre-hybridization buffer (PreHB, 0.1% Tween-20, 35% formamide, 2xSSC) with dilution factors of 1:1000 and 1:2000, respectively.

#### 5. Silanization and PDL-coating of coverslips

50mm x 64mm glass coverslips (CellPath, SAF-5064-02A) were incubated in 18.5% HCl in methanol for 30 min at room temperature (RT) followed by 4 washes with MiliQ water. Then coverslips were incubated in 70% ethanol for 5 min at RT followed by drying at 60°C for 30 min. Next, coverslips were incubated in a solution containing 0.1% triethylamine and 0.2% allyl trichlorosilane in chloroform for 30 min at RT. Coverslips were then washed with chloroform followed by washing with 100% ethanol. Then coverslips were dried at 60°C for 30 min. Coverslips were stacked on top of each other with a working solution of Poly-D-Lysine (PDL, SIgmaAldrich, A-003-M) (1 mg PDL/ml + RNase inhibitor) in between coverslips and incubated for ∼3 hours. Next, coverslips were dried at 60°C for 30 min. Finally, silanized and PDL-coated coverslips were stored at −20°C until use (good for 6 months).

#### 6. MERFISH+ sample preparation

Human postmortem basal ganglia samples in OCT block were sectioned (16 µm thick) using cryostat (Leica CM1850) at −20°C and mounted onto PDL-coated pre-silanized coverslips as aforedescribed. The mounted tissue sections were air dried at room temperature for 5 minutes followed by fixation with 4% paraformaldehyde (PFA) in PBS for 15 min. The PFA-fixed samples were stored in a 10% glucose solution in PBS (with RNase inhibitor, 1:1000) at −80°C for future use. The PFA-fixed frozen samples were thawed and incubated with 70% (v/v) ethanol for 1 hour at room temperature. The samples were then incubated with 5% SDS (in 2xSSC) for 10 minutes at room temperature followed by three washes with 2xSSC. Samples were incubated with PreHB for 15 minutes at room temperature. The coverslip containing the sample was transferred to a new Petri dish and 200 µl of encoding probe hybridization buffer containing 200 µg of encoding probes were added onto the sample. The encoding probe hybridization buffer consisted of 50% (v/v) formamide in 2xSSC and 10% (wt/v) dextran sulfate, with the RNase inhibitor added. Parafilm was placed gently on the top of the sample to prevent evaporation of the encoding probe hybridization buffer (10% dextran sulfate, 50% formamide, 2xSSC) and incubated in humidified incubator for 18-24 hours at 47°C. After encoding probe hybridization, samples were washed with PreHB twice 15 minutes each at room temperature. Next, 4% polyacrylamide gel embedding of samples was performed to anchor RNA molecules at place. Samples were then post-fixed with 4% PFA for 10 minutes at room temperature. Samples were incubated with digestion buffer containing 2% SDS in 2x SSC and 2% proteinase K overnight at 37°C. To quench sample autofluorescence, samples (still in digestion buffer) were incubated in Vizgen photobleacher for at least 3 hours. Finally, samples were washed with 2x SSC three times and proceeded to image acquisition.

#### 7. Microscope setup for MERFISH+ imaging

MERFISH+ image acquisition was performed with a custom-built system consisting of a custom-built microscope and microfluidics systems (as described in ^44^). The microscope was built with an ASI microscope body coupled with Nikon CFI Plan Apo 60X Lambda D oil immersion objective. Custom software was used to control system components based on https://github.com/ZhuangLab/storm-control (RRID: SCR_027698) and also see the MERFISH+ reference ^44^

### MERFISH+ data acquisition

MERFISH+ data acquisition consisted of 10 sequential rounds of hybridization and imaging. Each of the hybridization round consisted of the following steps-(i) flow the adaptor probes (1:1000 dilution with PreHB) specific to each round and incubate for 5400 sec (ii) wash with formamide wash buffer (30% v/v formamide in 2xSSC + 0.1% Tween-20) (iii) flow the readout probes (1:2000 dilution with PreHB) and incubate for 5400 sec (iv) wash with formamide wash buffer (30% v/v formamide in 2xSSC+ 0.1% Tween-20) (v) flow 2xSSC (vi) flow imaging buffer. The composition of the imaging buffer was as previously described ^43^. After each round of hybridization, z-stack (10 µm) images were acquired with four channels (750 nm, 647 nm and 560 nm for RNA molecules; 405 nm for DAPI) at 0.4µm step size (25 frames per field of view/channel, total frames=100/field of view). After image acquisition of a round, fluorescence signals were removed through flowing stripping buffer (80% formamide in 0.8xSSC+0.1% Tween-20) and 2xSSC before proceeding to next round of hybridization and imaging.

### Bioinformatics

#### Stereo-seq data processing

We processed the Stereo-seq data using the standard analysis workflow software (https://github.com/STOmics/SAW, RRID:SCR_025001) version 8.3 with the following parameters: *–kit-version “Stereo-seq T FF V1.2”, – sequencing-type “PE100_50+100”.* The pipeline takes three files as an input: (1) paired-end FASTQ files, (2) mask file and (3) mosaic image of nuclei-staining. The first read in the *fastq* files contains coordinate ID (CID) - spatial location identifier barcode sequence that is unique to each DNA nano ball on the stereo-seq chip, and the Molecular ID (MID) - artificially synthesized mRNA-specific sequence that helps differentiate the number of reads contributed by mRNA expression level due to amplification. The CIDs were first aligned and matched with the values in the mask file which contains the mapping of the CID barcodes to the actual spatial coordinates in the tissue. This allows the identification of the spatial location of the mRNAs within the tissue section. Reads that have MID sequence with more than one N bases or more than one bases with quality less than Q10 were excluded. The second read in the fastq file contains the actual sequence of the captured mRNA. These were aligned to a reference genome using the STAR aligner. The reference genome and transcript annotation were done using the human *GRCh38.dna* primary sequence assembly and *GRCh38.113* GTF files downloaded from Ensembl (RRID:SCR_002344). The cell segmentation was performed by applying a combination of the watershed algorithm, the contrast limited adaptive histogram equalization (CLAHE) algorithm and the BCDU-Net mode on the ssDNA nuclei-staining images. Finally, the cell segmentation mask and the count of the transcripts were combined to generate a cell-by-gene count matrix.

### MERFISH image processing

MERFISH image analysis consists of 4 major steps: (1) detecting individual fluorescence spots; (2) correcting the drift between the images across the different hybridization rounds; (3) decoding the gene identity of fluorescence spots by matching their on-off intensity pattern with the binary words in the codebook; (4) segmenting the mask of individual cells. A detailed overview of each step is as follows.

#### 1. Spot detection

we applied flat-field correction to the raw transcript images in the Cy3, Cy5 and 750 channels to minimize vignetting artifacts. Next, we deconvolved the images by applying the Wiener’s algorithm as implemented in the *sdeconv* package using a point spread function (PSF) estimated from the raw transcript images. Then, we normalized the resulting images and detected spots with gaussian peaks.

#### 2. Drift correction

To estimate the drift between hybridization rounds, we detected image features corresponding to local minima and local maxima intensity points within the DAPI channel using the same procedure as in the spot detection step. These features were then used to register the images from different hybridization rounds using phase cross-correlation.

#### 3. Decoding

We decoded the detected fluorescent spots as described previously ^42^. Briefly, we matched each spot in an image to spots that are within a set distance threshold in other images across the different hybridization rounds. We then compared the intensity pattern formed by the colocalizing spots to the binary words in the codebook in order to decode their gene identity. Finally, we filtered the molecules based on their correlation to the point spread function and the distance of their normalized brightness from the matched binary word.

#### 4. Cell segmentation

First, we deconvolved the flat-field corrected DAPI images using the Wiener algorithm, with the parameter beta set to 0.01, in order to improve the contrast of the nuclei boundaries. Then for each z-plane, we performed cell segmentation using the Cellpose ‘nuclei’ model ^79^ with the parameters: ***diameter*** = 23, ***flow_threshold*** = −10 and ***cellprob_threshold*** = −10. Next, we stitched the segmented images to get a 3D reconstruction of the cell masks. We then applied the masks to the decoded molecule in order to assign them to cells based on their spatial position. Finally, we generated a cell-by-gene count matrix by aggregating the transcript count of each gene in every cell. In addition, we generated cell metadata that included spatial information such as the XYZ location and volume of the imaged cells.

### Cell type clustering and cell type annotation

#### 1. Pre-processing

We preprocessed the cell by gene count matrix as follows. First, cells with total transcript counts greater than 99 percentile or less 1 percentile were filtered out. Second, cells with very large (above 98 percentile) and very small (below 2 percentile) volume were removed. Such cells arise from segmentation artifacts and usually represent incomplete or doublet cells (two different cells segmented as one). The count matrix was then normalized by volume of cells, in order to account for the effect of cell size variation on the expression level of genes. The matrix was again normalized by the total count of transcripts in the cells in order to reduce noise arising from the difference in the relative expression of genes between cells. Finally, the data was log-normalized to ensure all genes had the same expression baseline and hence minimized the risk of masking low abundant genes.

#### 2. Clustering and annotation

We performed clustering analysis using classical single-cell clustering pipeline from the Scanpy package^80^. First, the graph representation of the count matrix was generated using the *neighbors* module, *scanpy.pp.neighbors(use_rep = ‘X’, n_neighbors = 15)*. Then, the communities formed in the graph were clustered using the Leiden clustering algorithm, *<u>scanpy.tl</u>.leiden(resolution = 1).* Finally, the UMAP embedding of the clustered data was obtained by projecting the high dimensional clustering into 2D space, *<u>scanpy.tl</u>.umap(resolution = 1, min_dist = 0.1).* The annotation of cell types was done by mapping the data to the Cross-species Basal ganglion cell type taxonomy from the Allen institute through cell_type_mapper (the python backend for MapMyCells, RRID:SCR_024672) using two precomputed supporting files shared through the BICAN consortium: *query_markers (a lookup table of marker genes for the taxonomy) and precomputed_stats (HDF5 file that defines the taxonomy)*.

We performed the subclustering analysis for astrocytes using only astrocytic genes in order to avoid artifacts from cross-talk or contamination from other cell types. To identify the genes, we applied the Wilcoxon rank sum method on the pre-processed count matrix as implemented in the *rank_genes_groups* function *from the Scanpy package*^80^. Next, we reduced the data to cells that were labeled as astrocytes at the Subclass level and to genes that were differentially expressed in them. Following this, we performed further quantity control to remove outlier cells with volume and transcript count below 1 percentile or greater than 99th percentile. Next, we did Principal Component Analysis (PCA) and harmony integration on the normalized count matrix using the functions *sc.pl.pca* and *sc.external.pp.harmony_intergrate* from Scanpy respectively. We then constructed the KNN graph based on the principal components from harmony integration using the parameters n_neighbors = 10 and n_pcs = 20. Lastly, we ran Leiden clustering with a resolution of 0.6.

### Spatial module analysis

We performed spatial module analysis with two different approaches, (1) cell-type–based community clustering from MERFISH+ data, and (2) whole-transcriptome spatial clustering from Stereo-seq data. Together, these analyses resolve both mesoscale anatomical territories and microscale compartmental organization within the human basal ganglia.

#### 1. Spatial module clustering based on cell type composition

Using cell type composition, we delineated both major anatomical regions of the basal ganglia such as Putamen, Caudate, NAC core, NAC shell and internal and external segment of the globus palladium; and the local compartments of the striatum including the Striosome and Matrix.

For each cell, we first searched neighboring cells that were within a radius r in the spatial coordinate of the tissue using the *KDTree* module from *SciPy* (RRID:SCR_008058) ^81^. Then we counted the number of cell types at the group level in the identified neighborhoods to construct the cell type composition matrix. The rows of the matrix correspond to individual cells and the columns are the cell types at group level. Then we generated a KNN graph based on the composition matrix using *<u>scanpy.pp</u>.neighbors*, and then we ran Leiden clustering using *<u>scanpy.pp</u>.leiden* to cluster the cells into spatial modules. The clustering was performed hierarchically; we first identified high level modules that corresponded to major anatomical regions such as the white matter and the Striatum. Then we iteratively subclustered each module to identify more finer regions. During each iteration, we tuned different parameters including the radius for searching spatially neighboring cells, the number of neighbors used for the KNN graph, and the resolution for Leiden clustering until the resulting modules captured known anatomical regions. The identified spatial modules corresponded well to anatomical regions defined in the Allen human Brain atlas^59^; however, the internal globus pallidus module partially overlapped with adjacent white matter tracts; its boundary was refined using the spatial location of GPi Shell neurons, which delineated the GPi border with high precision.

To identify spatial modules corresponding to the striosome and matrix compartment, we followed similar but slightly different procedure to that used in global spatial modules analysis. The main differences were that we only used MSN cell types when building the cell type composition matrix and we only included the cells in the striatum for the analysis as the Striosome and Matrix compartments exist only in this part of the Basal ganglia. In detail, we first built a K-nearest neighbor graph based on the spatial location of MSN subtypes using the scikit-learn package^82^, then we queried the graph to identify 50 nearest MSNs for each cell in the striatum. Next, as explained in ^14^, we defined the composition matrix as distance weighted frequency count of neighboring MSN subtypes. This matrix was then L2 normalized and clustered using the Leiden algorithm from the Scanpy package. The same analysis approach was used in human and mouse datasets.

#### 2. Whole transcriptome based spatial module analysis

To do spatial module analysis using gene expression, we integrated the expression values with the spatial information of cells. First, we calculated the spatial distance of each cell to neighboring cells located within a radius r using the function *radius_neighbors_graph* from *scikit-learn*^82^. Next, we took the negative exponential of the distance values and normalized them by the total distance to get a gaussian distribution. The resulting values were then multiplied with the gene expression matrix to get spatially weighted average gene expression for each cell. Next, using the Scanpy package^80^, we identified the top 5000 highly variable genes, which we then used for PCA and KNN graph computation. Finally, we did Leiden clustering to identify the spatial modules. Similar to the spatial modules analysis based on cell type composition, we did the clustering hierarchically - starting from high level to finer spatial modules. And the parameters were tuned at each hierarchy until the modules matched the anatomical regions.

### Cell type enrichment analysis

#### Cell type enrichment analyses across spatial anatomical modules

The enrichment analysis was done following the method described in Zhang *et al*. ^14^. We first computed the confusion matrix between spatial modules and cell types at the group level using the *pivot_table* function from the pandas package. Next, for each cell type, we calculated its expected average density in every spatial module as the product of the fraction of cells that belonged to the cell type and the fraction of cells found in the spatial modules. Then the enrichment score of a cell type in a spatial module was computed as the ratio of its cell density and expected average cell density in the spatial module. To quantify the significance of the result, we used a permutation test where we shuffled cell type labels and recomputed the enrichment score. We repeated this step 1000 times to generate a null distribution of the enrichment score. Then the p-value was computed as the fraction of values in the distribution that are at least as large as the observed enrichment score. Finally, we corrected the p-values for multiple testing using the Bonferroni method.

#### Cell type enrichment analyses across the matrix and striosome compartments

We quantified the enrichment of cell types within the Matrix and Striosome compartments using the Cochran-Mantel-Haenszel statistics^83^. For each cell type, we built contingency tables that summarize the number of cells in the compartments from each individual sample. Then, we computed the common odds-ratio as:

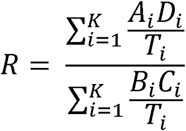

Where,

K : is the total number of samples,

*A_i_* : is the number of cells in sample *i* that belongs to a cell type of interest and are within the brain region of interest.

*B_i_* : is the number of cells in sample *i* that belongs to a cell type of interest but are not within the brain region of interest.

*C_i_* : is the number of cells in sample *i* that does not belong to a cell type of interest but are within the brain region of interest.

*D*_i_: is the number of cells in sample *i* that does not belong to a cell type of interest and are not within the brain region of interest.

*T*_i_ : is the total number of cells in the data.

Next, we quantified the significance of the result using the permutation test, as described above.

### Cell type organization and gene expression gradients with respect to striosome borders

We first identified the contours for striosome borders from the spatial map of the brain section using the function findcontours from the Opencv package. Then we used the KDTree module from *SciPy*^81^ to find the distance of each cell from the contour of the nearest striosome. We set the distance of cells within the striosome compartment as positive and negative for those that are outside. Then we binned the distance into bins of size 50 microns and computed the mean abundance of celltype within each bin. When doing the calculation, we only considered cells that are 200 microns away from the striosome border and 500 microns into the center of the striosome. We also removed cell types that were very sparse - with less than 1 % abundance near the striosome.

To quantify the gene expression gradient with respect to the border striosomes, we computed the mean log1p gene expression of cells that are within the binned distances. Then, we performed *lowess regression* using the statsmodel package to quantify the dependence of the mean gene expression on the distance of cells from the striosomes. We computed the significance of the estimated values using a permutation test. More specifically, we randomized the distance values and recomputed the regression and, then we obtained the explained variance for the estimated values. We repeated the permutation 1000 times to generate a null distribution, the p-value was then calculated as the fraction of the explained variance that were at least as greater as the observed explained variance.

### Analysis of gene expression gradients in the striatum using stereo-seq data

To quantify the gene expression gradient along the ventromedial to dorsolateral (VM-DL) axis, we used the internal capsule as our reference since it is located approximately along this axis. We first rotated the brain sections so that the internal capsule aligns with the horizontal axis. Then we digitized the rotated coordinate values using the combination of the functions *digitize and histogram* from the NumPy package (RRID:SCR_008633). Next, we grouped the cells located within the same bin (digitized coordinates) and took the average of their spatially weighted gene expression values. Following this, we quantified the expression gradient of each gene by calculating the correlation between the bin average gene expression and the axis coordinates using the ***pearsonr*** function from *SciPy.* Finally, we integrated the results from all stereo-seq samples by computing the mean and std of the correlation coefficients. We only kept genes whose standard deviation is below 0.2 and reported their mean correlation coefficient as final value.

### Gene Ontology enrichment analysis for spatial modules

We performed gene ontology enrichment analysis using the *enricher* function from the *gget* package using the “ontology” database. For each spatial module, we first identified the marker genes that were differentially expressed in each spatial module. Then we obtained the top 10 ontology terms in relation to these genes by running the *enricher* function. Next, we mapped each term to their parent (broad) categories by parsing the directed acyclic graph (DAG) built from the basic GO ontology file - ‘go-basic.obo’ using the *GODag* module from *goatools* package. Finally, we computed the frequency of the parent terms in each module and calculated the proportion of the frequencies across the regions to quantify the relative abundance across the regions.

### Density-based clustering of subcellular transcript distributions

For each cell type, we first selected 100 cells whose differential gene expression correlated most with the average (pseudobulk) differential gene expression of the cell type. Then, we quantified the enrichment of every gene within the intracellular space of the cells as the fold change between transcript counts inside and outside the cell’s segmentation mask. We excluded genes from further analysis if their enrichment was less than 3-fold or if they were not differentially expressed. Next, we computed the median distance between neighboring transcripts in both the somatic and putative distal neuritic spaces using the KDTree function from the SciPy package^81^. Following this, we performed density-based clustering to identify the somatic transcripts corresponding to each cell. To achieve this, we used the DBSCAN function from the scikit-learn package (RRID:SCR_002577) ^82^ with the maximum distance parameter set to the average of the median transcript distances in somatic and putative distal neuritic spaces. Next, we classified the non-somatic transcripts as neuritic if they are within 20 microns from the nearest somatic transcript.

### Computing the volume of cells using somatic transcripts

To quantify the volume of a cell, we generated a tetrahedral mesh from the point cloud formed by the cell’s somatic transcripts. For this, we performed 3D Delaunay triangulation using the function *delaunay_3d* from the *pyvista* package ^84^ with the parameter alpha set to 3 times the median distance between neighboring transcripts. We then triangulated the mesh and computed its volume using the *triangulate* function. Following this procedure, we quantified the volume of 100 cells sampled from each cell type. Finally, we took the median of the cell’s volume to get a representative value for each cell type. We filtered out cells that had artifacts in the labeling of their somatic transcripts. Such artifacts arise when cells are surrounded by density transcript from nearby cells or their processes.

### Quantifying the projection of neurons using distribution of subcellular transcripts

As the MERFISH imaging data is acquired on a field of view by field of view basis, the decoded transcript information is organized separately for each FOV. For this reason, we performed the quantification of a neuron projection to a target region by aggregating the information within each FOV. For every gene in each FOV, we first computed the count of transcripts that are outside a cell’s segmentation mask. Then we took the log of the counts and normalized them across genes and all FOVs. Next, we computed the differential gene expression between *STRd D1 Matrix* and *STRd D2 matrix MSNs*. The differential expression was quantified as a difference-in-difference: the deviation of *STRd D1 Matrix MSN’s* pseudobulk expression from the across-cell-type mean, minus the deviation of *STRd D2 matrix MSN’s* pseudobulk expression from that mean. Finally, we quantified the projection strength as the dot product between log count of extracellular transcripts and the differential gene expression scores.

### Identifying mouse homologues of human basal ganglia cell types

To identify the mouse homologues of the cell types detected in our human Basal ganglia MERFISH dataset, we used the imputed whole mouse brain MERFISH spatial transcriptomic dataset from an earlier publication^10^. First, we manually insepcted the mouse MERFISH data to identify tissue sections that contain brain regions corresponding to those covered in the human samples. We selected the sections of C57BL6J-638850.39 - 50, which show the caudoputamen and globus pallidus regions. Then we used the parcellation_division label “STR” and “PAL” in the dataset to select all cells located in the mouse striatum and pallidum. Next, we filtered the data by removing low abundance cell types; cell types with less than 150 cell counts. We then mapped the mouse genes to their human orthologs using the mouse human homologous gene table from the Mouse Genome informatics (MGI) database. Next, we identified the genes shared between the mouse and human MERFISH dataset and reduced the gene expression matrix in both mouse and human to the common genes. Following this, we computed the pseudobulk gene expression of each cell type at the supertype level for the mouse and at the group level for the human. Then we performed the Pearson correlation between the mouse and human expression values. Through this approach, we identified the mouse homologs of the human D1 Matrix and Striosome MSNs. Both D2 Matrix and Striosome MSNs were initially mapped to 0980 STR D2 Gaba_4 mouse cell type; we differentiated the two cell types after subclustering this supertype as 0277 STR D2 Gaba_4_A* which corresponds to STRd striosomal MSN and 0277 STR D2 Gaba_4_B* which corresponds to the STRd matrix MSN.

## References

1. Graybiel, A.M. (2000). The basal ganglia. Curr Biol 10, R509–511. 10.1016/s0960-9822(00)00593-5.

2. Mendelsohn, A.I., Nikoobakht, L., Bikoff, J.B., and Costa, R.M. (2025). Segregated basal ganglia output pathways correspond to genetically divergent neuronal subclasses. Cell Rep 44, 115454. 10.1016/j.celrep.2025.115454.

3. Lanciego, J.L., Luquin, N., and Obeso, J.A. (2012). Functional neuroanatomy of the basal ganglia. Cold Spring Harb Perspect Med 2, a009621. 10.1101/cshperspect.a009621.

4. He, J., Kleyman, M., Chen, J., Alikaya, A., Rothenhoefer, K.M., Ozturk, B.E., Wirthlin, M., Bostan, A.C., Fish, K., Byrne, L.C., et al. (2021). Transcriptional and anatomical diversity of medium spiny neurons in the primate striatum. Curr Biol 31, 5473–5486 e5476. 10.1016/j.cub.2021.10.015.

5. Krienen, F.M., Levandowski, K.M., Zaniewski, H., Del Rosario, R.C.H., Schroeder, M.E., Goldman, M., Wienisch, M., Lutservitz, A., Beja-Glasser, V.F., Chen, C., et al. (2023). A marmoset brain cell census reveals regional specialization of cellular identities. Sci Adv 9, eadk3986. 10.1126/sciadv.adk3986.

6. Calabresi, P., Picconi, B., Tozzi, A., Ghiglieri, V., and Di Filippo, M. (2014). Direct and indirect pathways of basal ganglia: a critical reappraisal. Nat Neurosci 17, 1022–1030. 10.1038/nn.3743.

7. Kerkhoff, W.G., and Stauffer, W.R. (2024). Basal ganglia: Uniting circuit logic between matrix and striosome. Curr Biol 34, R1149–R1152. 10.1016/j.cub.2024.10.010.

8. Lazaridis, I., Crittenden, J.R., Ahn, G., Hirokane, K., Wickersham, I.R., Yoshida, T., Mahar, A., Skara, V., Loftus, J.H., Parvataneni, K., et al. (2024). Striosomes control dopamine via dual pathways paralleling canonical basal ganglia circuits. Curr Biol 34, 5263–5283 e5268. 10.1016/j.cub.2024.09.070.

9. Siletti, K., Hodge, R., Mossi Albiach, A., Lee, K.W., Ding, S.L., Hu, L., Lonnerberg, P., Bakken, T., Casper, T., Clark, M., et al. (2023). Transcriptomic diversity of cell types across the adult human brain. Science 382, eadd7046. 10.1126/science.add7046.

10. Yao, Z., van Velthoven, C.T.J., Kunst, M., Zhang, M., McMillen, D., Lee, C., Jung, W., Goldy, J., Abdelhak, A., Aitken, M., et al. (2023). A high-resolution transcriptomic and spatial atlas of cell types in the whole mouse brain. Nature 624, 317–332. 10.1038/s41586-023-06812-z.

11. Garma, L.D., Harder, L., Barba-Reyes, J.M., Marco Salas, S., Diez-Salguero, M., Nilsson, M., Serrano-Pozo, A., Hyman, B.T., and Munoz-Manchado, A.B. (2024). Interneuron diversity in the human dorsal striatum. Nat Commun 15, 6164. 10.1038/s41467-024-50414-w.

12. Roseberry, T.K., Lee, A.M., Lalive, A.L., Wilbrecht, L., Bonci, A., and Kreitzer, A.C. (2016). Cell-Type-Specific Control of Brainstem Locomotor Circuits by Basal Ganglia. Cell 164, 526–537. 10.1016/j.cell.2015.12.037.

13. Mandali, A., Srinivasa Chakravarthy, V., and Moustafa, A.A. (2018). The Molecular, Cellular, and Systems-Level Structure of the Basal Ganglia. In Computational Neuroscience Models of the Basal Ganglia, V.S. Chakravarthy, and A.A. Moustafa, eds. (Springer Singapore), pp. 5–19. 10.1007/978-981-10-8494-2_2.

14. Zhang, M., Pan, X., Jung, W., Halpern, A.R., Eichhorn, S.W., Lei, Z., Cohen, L., Smith, K.A., Tasic, B., Yao, Z., et al. (2023). Molecularly defined and spatially resolved cell atlas of the whole mouse brain. Nature 624, 343–354. 10.1038/s41586-023-06808-9.

15. Tran, M.N., Maynard, K.R., Spangler, A., Huuki, L.A., Montgomery, K.D., Sadashivaiah, V., Tippani, M., Barry, B.K., Hancock, D.B., Hicks, S.C., et al. (2021). Single-nucleus transcriptome analysis reveals cell-type-specific molecular signatures across reward circuitry in the human brain. Neuron 109, 3088–3103 e3085. 10.1016/j.neuron.2021.09.001.

16. Kalanithi, P.S., Zheng, W., Kataoka, Y., DiFiglia, M., Grantz, H., Saper, C.B., Schwartz, M.L., Leckman, J.F., and Vaccarino, F.M. (2005). Altered parvalbumin-positive neuron distribution in basal ganglia of individuals with Tourette syndrome. Proc Natl Acad Sci U S A 102, 13307–13312. 10.1073/pnas.0502624102.

17. Courtney, C.D., Pamukcu, A., and Chan, C.S. (2023). Cell and circuit complexity of the external globus pallidus. Nat Neurosci 26, 1147–1159. 10.1038/s41593-023-01368-7.

18. Sandberg, M., Flandin, P., Silberberg, S., Su-Feher, L., Price, J.D., Hu, J.S., Kim, C., Visel, A., Nord, A.S., and Rubenstein, J.L.R. (2016). Transcriptional Networks Controlled by NKX2-1 in the Development of Forebrain GABAergic Neurons. Neuron 91, 1260–1275. 10.1016/j.neuron.2016.08.020.

19. Saunders, A., Macosko, E.Z., Wysoker, A., Goldman, M., Krienen, F.M., de Rivera, H., Bien, E., Baum, M., Bortolin, L., Wang, S., et al. (2018). Molecular Diversity and Specializations among the Cells of the Adult Mouse Brain. Cell 174, 1015–1030 e1016. 10.1016/j.cell.2018.07.028.

20. Baker, M., Kang, S., Hong, S.I., Song, M., Yang, M.A., Peyton, L., Essa, H., Lee, S.W., and Choi, D.S. (2023). External globus pallidus input to the dorsal striatum regulates habitual seeking behavior in male mice. Nat Commun 14, 4085. 10.1038/s41467-023-39545-8.

21. Gast, R., Gong, R., Schmidt, H., Meijer, H.G.E., and Knosche, T.R. (2021). On the Role of Arkypallidal and Prototypical Neurons for Phase Transitions in the External Pallidum. J Neurosci 41, 6673–6683. 10.1523/JNEUROSCI.0094-21.2021.

22. Chen, W.T., Lu, A., Craessaerts, K., Pavie, B., Sala Frigerio, C., Corthout, N., Qian, X., Lalakova, J., Kuhnemund, M., Voytyuk, I., et al. (2020). Spatial Transcriptomics and In Situ Sequencing to Study Alzheimer’s Disease. Cell 182, 976–991 e919. 10.1016/j.cell.2020.06.038.

23. Piwecka, M., Rajewsky, N., and Rybak-Wolf, A. (2023). Single-cell and spatial transcriptomics: deciphering brain complexity in health and disease. Nat Rev Neurol 19, 346–362. 10.1038/s41582-023-00809-y.

24. Rao, A., Barkley, D., Franca, G.S., and Yanai, I. (2021). Exploring tissue architecture using spatial transcriptomics. Nature 596, 211–220. 10.1038/s41586-021-03634-9.

25. Ortiz, C., Carlen, M., and Meletis, K. (2021). Spatial Transcriptomics: Molecular Maps of the Mammalian Brain. Annu Rev Neurosci 44, 547–562. 10.1146/annurev-neuro-100520-082639.

26. Cheng, Y., Dang, S., Zhang, Y., Chen, Y., Yu, R., Liu, M., Jin, S., Han, A., Katz, S., and Wang, S. (2025). Sequencing-free whole-genome spatial transcriptomics at single-molecule resolution. Cell. 10.1016/j.cell.2025.09.006.

27. Lei, Y., Liu, Y., Wang, M., Yuan, N., Hou, Y., Ding, L., Zhu, Z., Wu, Z., Li, C., Zheng, M., et al. (2025). Single-cell spatial transcriptome atlas and whole-brain connectivity of the macaque claustrum. Cell. 10.1016/j.cell.2025.02.037.

28. Del Rey, N.L., and Garcia-Cabezas, M.A. (2023). Cytology, architecture, development, and connections of the primate striatum: Hints for human pathology. Neurobiol Dis 176, 105945. 10.1016/j.nbd.2022.105945.

29. Chen, K.H., Boettiger, A.N., Moffitt, J.R., Wang, S., and Zhuang, X. (2015). RNA imaging. Spatially resolved, highly multiplexed RNA profiling in single cells. Science 348, aaa6090. 10.1126/science.aaa6090.

30. Moffitt, J.R., and Zhuang, X. (2016). RNA Imaging with Multiplexed Error-Robust Fluorescence In Situ Hybridization (MERFISH). Methods Enzymol 572, 1–49. 10.1016/bs.mie.2016.03.020.

31. Moffitt, J.R., Bambah-Mukku, D., Eichhorn, S.W., Vaughn, E., Shekhar, K., Perez, J.D., Rubinstein, N.D., Hao, J., Regev, A., Dulac, C., and Zhuang, X. (2018). Molecular, spatial, and functional single-cell profiling of the hypothalamic preoptic region. Science 362. 10.1126/science.aau5324.

32. Moffitt, J.R., Hao, J., Bambah-Mukku, D., Lu, T., Dulac, C., and Zhuang, X. (2016). High-performance multiplexed fluorescence in situ hybridization in culture and tissue with matrix imprinting and clearing. Proc Natl Acad Sci U S A 113, 14456–14461. 10.1073/pnas.1617699113.

33. Zhuang, X. (2021). Spatially resolved single-cell genomics and transcriptomics by imaging. Nat Methods 18, 18–22. 10.1038/s41592-020-01037-8.

34. Chen, A., Liao, S., Cheng, M., Ma, K., Wu, L., Lai, Y., Qiu, X., Yang, J., Xu, J., Hao, S., et al. (2022). Spatiotemporal transcriptomic atlas of mouse organogenesis using DNA nanoball-patterned arrays. Cell 185, 1777–1792 e1721. 10.1016/j.cell.2022.04.003.

35. You, Y., Fu, Y., Li, L., Zhang, Z., Jia, S., Lu, S., Ren, W., Liu, Y., Xu, Y., Liu, X., et al. (2024). Systematic comparison of sequencing-based spatial transcriptomic methods. Nat Methods 21, 1743–1754. 10.1038/s41592-024-02325-3.

36. Chen, A., Sun, Y., Lei, Y., Li, C., Liao, S., Meng, J., Bai, Y., Liu, Z., Liang, Z., Zhu, Z., et al. (2023). Single-cell spatial transcriptome reveals cell-type organization in the macaque cortex. Cell 186, 3726–3743 e3724. 10.1016/j.cell.2023.06.009.

37. Hao, S., Zhu, X., Huang, Z., Yang, Q., Liu, H., Wu, Y., Zhan, Y., Dong, Y., Li, C., Wang, H., et al. (2024). Cross-species single-cell spatial transcriptomic atlases of the cerebellar cortex. Science 385, eado3927. 10.1126/science.ado3927.

38. Lei, Y., Liu, Y., Wang, M., Yuan, N., Hou, Y., Ding, L., Zhu, Z., Wu, Z., Li, C., Zheng, M., et al. (2025). Single-cell spatial transcriptome atlas and whole-brain connectivity of the macaque claustrum. Cell 188, 3863–3881 e3825. 10.1016/j.cell.2025.02.037.

39. Fang, R., Xia, C., Close, J.L., Zhang, M., He, J., Huang, Z., Halpern, A.R., Long, B., Miller, J.A., Lein, E.S., and Zhuang, X. (2022). Conservation and divergence of cortical cell organization in human and mouse revealed by MERFISH. Science 377, 56–62. 10.1126/science.abm1741.

40. Xie, F., Jain, S., Xu, R., Butrus, S., Tan, Z., Xu, X., Shekhar, K., and Zipursky, S.L. (2025). Spatial profiling of the interplay between cell type- and vision-dependent transcriptomic programs in the visual cortex. Proc Natl Acad Sci U S A 122, e2421022122. 10.1073/pnas.2421022122.

41. Johnston, K.G., Berackey, B.T., Tran, K.M., Gelber, A., Yu, Z., MacGregor, G.R., Mukamel, E.A., Tan, Z., Green, K.N., and Xu, X. (2025). Single-cell spatial transcriptomics reveals distinct patterns of dysregulation in non-neuronal and neuronal cells induced by the Trem2(R47H) Alzheimer’s risk gene mutation. Mol Psychiatry 30, 461–477. 10.1038/s41380-024-02651-0.

42. Maimon, R., Chillon-Marinas, C., Vazquez-Sanchez, S., Kern, C., Jenie, K., Malukhina, K., Moore, S., Cui, J., Goginashvili, A., Moghadami, S., et al. (2024). Re-activation of neurogenic niches in aging brain. bioRxiv. 10.1101/2024.01.27.575940.

43. Su, J.H., Zheng, P., Kinrot, S.S., Bintu, B., and Zhuang, X. (2020). Genome-Scale Imaging of the 3D Organization and Transcriptional Activity of Chromatin. Cell 182, 1641–1659 e1626. 10.1016/j.cell.2020.07.032.

44. Kern, C., Zhang, Q., Lu, Y., Eschbach, J., Zeng, Z., Farah, E.N., Tai, C.-Y., Yang, K., Jenie, I., Yao, F., et al. (2025). MERFISH+, a large-scale, multi-omics spatial technology resolves the molecular holograms of the 3D human developing heart bioRxiv. 10.1101/2025.11.02.686137.

45. Hewitt, M.N., Turner, M.A., Johansen, N., McMillen, D.A., Dan, S., DeBerardine, M., Ruiz, A., Huang, M., Quon, J., Fu, Y., et al. (2025). A cross-species spatial transcriptomic atlas of the human and non-human primate basal ganglia. bioRxiv, 2025.2011.2022.688128. 10.1101/2025.11.22.688128.

46. Surmeier, D.J., Ding, J., Day, M., Wang, Z., and Shen, W. (2007). D1 and D2 dopamine-receptor modulation of striatal glutamatergic signaling in striatal medium spiny neurons. Trends Neurosci 30, 228–235. 10.1016/j.tins.2007.03.008.

47. Gagnon, D., Petryszyn, S., Sanchez, M.G., Bories, C., Beaulieu, J.M., De Koninck, Y., Parent, A., and Parent, M. (2017). Striatal Neurons Expressing D(1) and D(2) Receptors are Morphologically Distinct and Differently Affected by Dopamine Denervation in Mice. Sci Rep 7, 41432. 10.1038/srep41432.

48. Perreault, M.L., Hasbi, A., Alijaniaram, M., Fan, T., Varghese, G., Fletcher, P.J., Seeman, P., O’Dowd, B.F., and George, S.R. (2010). The dopamine D1-D2 receptor heteromer localizes in dynorphin/enkephalin neurons: increased high affinity state following amphetamine and in schizophrenia. J Biol Chem 285, 36625–36634. 10.1074/jbc.M110.159954.

49. Salcedo, C., Wagner, A., Andersen, J.V., Vinten, K.T., Waagepetersen, H.S., Schousboe, A., Freude, K.K., and Aldana, B.I. (2021). Downregulation of GABA Transporter 3 (GAT3) is Associated with Deficient Oxidative GABA Metabolism in Human Induced Pluripotent Stem Cell-Derived Astrocytes in Alzheimer’s Disease. Neurochem Res 46, 2676–2686. 10.1007/s11064-021-03276-3.

50. Yadav, R., Han, G.W., and Gati, C. (2025). Molecular basis of human GABA transporter 3 inhibition. Nat Commun 16, 3830. 10.1038/s41467-025-59066-w.

51. Carrasco, A., Oorschot, D.E., Barzaghi, P., and Wickens, J.R. (2022). Three-Dimensional Spatial Analyses of Cholinergic Neuronal Distributions Across The Mouse Septum, Nucleus Basalis, Globus Pallidus, Nucleus Accumbens, and Caudate-Putamen. Neuroinformatics 20, 1121–1136. 10.1007/s12021-022-09588-1.

52. Kozlov, A., Blazquez-Llorca, L., Benavides-Piccione, R., Kastanauskaite, A., Rojo, A.I., Munoz, A., Cuadrado, A., DeFelipe, J., and Grillner, S. (2025). Mouse and human striatal projection neurons compared - somatodendritic arbor, spines and in silico analyses. PLoS Comput Biol 21, e1013569. 10.1371/journal.pcbi.1013569.

53. Singh-Bains, M.K., Tippett, L.J., Hogg, V.M., Synek, B.J., Roxburgh, R.H., Waldvogel, H.J., and Faull, R.L. (2016). Globus pallidus degeneration and clinicopathological features of Huntington disease. Ann Neurol 80, 185–201. 10.1002/ana.24694.

54. Crittenden, J.R., and Graybiel, A.M. (2011). Basal Ganglia disorders associated with imbalances in the striatal striosome and matrix compartments. Front Neuroanat 5, 59. 10.3389/fnana.2011.00059.

55. Tran, T.S., Kolodkin, A.L., and Bharadwaj, R. (2007). Semaphorin regulation of cellular morphology. Annu Rev Cell Dev Biol 23, 263–292. 10.1146/annurev.cellbio.22.010605.093554.

56. Alto, L.T., and Terman, J.R. (2017). Semaphorins and their Signaling Mechanisms. Methods Mol Biol 1493, 1–25. 10.1007/978-1-4939-6448-2_1.

57. Koropouli, E., and Kolodkin, A.L. (2014). Semaphorins and the dynamic regulation of synapse assembly, refinement, and function. Curr Opin Neurobiol 27, 1–7. 10.1016/j.conb.2014.02.005.

58. Farah, E.N., Hu, R.K., Kern, C., Zhang, Q., Lu, T.Y., Ma, Q., Tran, S., Zhang, B., Carlin, D., Monell, A., et al. (2024). Spatially organized cellular communities form the developing human heart. Nature 627, 854–864. 10.1038/s41586-024-07171-z.

59. Ding, S.L., Royall, J.J., Sunkin, S.M., Ng, L., Facer, B.A., Lesnar, P., Guillozet-Bongaarts, A., McMurray, B., Szafer, A., Dolbeare, T.A., et al. (2016). Comprehensive cellular-resolution atlas of the adult human brain. J Comp Neurol 524, 3127–3481. 10.1002/cne.24080.

60. Matsushima, A., Pineda, S.S., Crittenden, J.R., Lee, H., Galani, K., Mantero, J., Tombaugh, G., Kellis, M., Heiman, M., and Graybiel, A.M. (2023). Transcriptional vulnerabilities of striatal neurons in human and rodent models of Huntington’s disease. Nat Commun 14, 282. 10.1038/s41467-022-35752-x.

61. Graybiel, A.M., and Matsushima, A. (2023). Striosomes and Matrisomes: Scaffolds for Dynamic Coupling of Volition and Action. Annu Rev Neurosci 46, 359–380. 10.1146/annurev-neuro-121522-025740.

62. Krienen, F.M., Goldman, M., Zhang, Q., R, C.H.D.R., Florio, M., Machold, R., Saunders, A., Levandowski, K., Zaniewski, H., Schuman, B., et al. (2020). Innovations present in the primate interneuron repertoire. Nature 586, 262–269. 10.1038/s41586-020-2781-z.

63. Schmitz, M.T., Sandoval, K., Chen, C.P., Mostajo-Radji, M.A., Seeley, W.W., Nowakowski, T.J., Ye, C.J., Paredes, M.F., and Pollen, A.A. (2022). The development and evolution of inhibitory neurons in primate cerebrum. Nature 603, 871–877. 10.1038/s41586-022-04510-w.

64. Corrigan, E.K., DeBerardine, M., Poddar, A., Turrero Garcia, M., de la, O.S., He, S., Sen, H., Duhne, M., Lindberg, S., Song, M., et al. (2025). Conservation and alteration of mammalian striatal interneurons. Nature 647, 187–193. 10.1038/s41586-025-09592-w.

65. Klug, J.R., Engelhardt, M.D., Cadman, C.N., Li, H., Smith, J.B., Ayala, S., Williams, E.W., Hoffman, H., and Jin, X. (2018). Differential inputs to striatal cholinergic and parvalbumin interneurons imply functional distinctions. Elife 7. 10.7554/eLife.35657.

66. Stanley, G., Gokce, O., Malenka, R.C., Sudhof, T.C., and Quake, S.R. (2020). Continuous and Discrete Neuron Types of the Adult Murine Striatum. Neuron 105, 688–699 e688. 10.1016/j.neuron.2019.11.004.

67. Grillner, S., and Robertson, B. (2016). The Basal Ganglia Over 500 Million Years. Curr Biol 26, R1088–R1100. 10.1016/j.cub.2016.06.041.

68. Foster, N.N., Barry, J., Korobkova, L., Garcia, L., Gao, L., Becerra, M., Sherafat, Y., Peng, B., Li, X., Choi, J.H., et al. (2021). The mouse cortico-basal ganglia-thalamic network. Nature 598, 188–194. 10.1038/s41586-021-03993-3.

69. Lai, H.M., Liu, A.K.L., Ng, H.H.M., Goldfinger, M.H., Chau, T.W., DeFelice, J., Tilley, B.S., Wong, W.M., Wu, W., and Gentleman, S.M. (2018). Next generation histology methods for three-dimensional imaging of fresh and archival human brain tissues. Nat Commun 9, 1066. 10.1038/s41467-018-03359-w.

70. Rosoklija, G.B., Petrushevski, V.M., Stankov, A., Dika, A., Jakovski, Z., Pavlovski, G., Davcheva, N., Lipkin, R., Schnieder, T., Scobie, K., et al. (2014). Reliable and durable Golgi staining of brain tissue from human autopsies and experimental animals. J Neurosci Methods 230, 20–29. 10.1016/j.jneumeth.2014.04.006.

71. Jang, H., Li, Y., Fung, A.A., Bagheri, P., Hoang, K., Skowronska-Krawczyk, D., Chen, X., Wu, J.Y., Bintu, B., and Shi, L. (2023). Super-resolution SRS microscopy with A-PoD. Nat Methods 20, 448–458. 10.1038/s41592-023-01779-1.

72. Walsh, C.L., Tafforeau, P., Wagner, W.L., Jafree, D.J., Bellier, A., Werlein, C., Kuhnel, M.P., Boller, E., Walker-Samuel, S., Robertus, J.L., et al. (2021). Imaging intact human organs with local resolution of cellular structures using hierarchical phase-contrast tomography. Nat Methods 18, 1532–1541. 10.1038/s41592-021-01317-x.

73. Nord, C.L., Kim, S.G., Callesen, M.B., Kvamme, T.L., Jensen, M., Pedersen, M.U., Thomsen, K.R., and Voon, V. (2019). The myeloarchitecture of impulsivity: premature responding in youth is associated with decreased myelination of ventral putamen. Neuropsychopharmacology 44, 1216–1223. 10.1038/s41386-019-0343-6.

74. Krishnan, S., Cler, G.J., Smith, H.J., Willis, H.E., Asaridou, S.S., Healy, M.P., Papp, D., and Watkins, K.E. (2022). Quantitative MRI reveals differences in striatal myelin in children with DLD. Elife 11. 10.7554/eLife.74242.

75. De Biase, L.M., Schuebel, K.E., Fusfeld, Z.H., Jair, K., Hawes, I.A., Cimbro, R., Zhang, H.Y., Liu, Q.R., Shen, H., Xi, Z.X., et al. (2017). Local Cues Establish and Maintain Region-Specific Phenotypes of Basal Ganglia Microglia. Neuron 95, 341–356 e346. 10.1016/j.neuron.2017.06.020.

76. Woolley, S.C., Rajan, R., Joshua, M., and Doupe, A.J. (2014). Emergence of context-dependent variability across a basal ganglia network. Neuron 82, 208–223. 10.1016/j.neuron.2014.01.039.

77. Niccolini, F., Foltynie, T., Reis Marques, T., Muhlert, N., Tziortzi, A.C., Searle, G.E., Natesan, S., Kapur, S., Rabiner, E.A., Gunn, R.N., et al. (2015). Loss of phosphodiesterase 10A expression is associated with progression and severity in Parkinson’s disease. Brain 138, 3003–3015. 10.1093/brain/awv219.

78. Beaumont, V., Zhong, S., Lin, H., Xu, W., Bradaia, A., Steidl, E., Gleyzes, M., Wadel, K., Buisson, B., Padovan-Neto, F.E., et al. (2016). Phosphodiesterase 10A Inhibition Improves Cortico-Basal Ganglia Function in Huntington’s Disease Models. Neuron 92, 1220–1237. 10.1016/j.neuron.2016.10.064.

79. Stringer, C., Wang, T., Michaelos, M., and Pachitariu, M. (2021). Cellpose: a generalist algorithm for cellular segmentation. Nat Methods 18, 100–106. 10.1038/s41592-020-01018-x.

80. Wolf, F.A., Angerer, P., and Theis, F.J. (2018). SCANPY: large-scale single-cell gene expression data analysis. Genome Biol 19, 15. 10.1186/s13059-017-1382-0.

81. Virtanen, P., Gommers, R., Oliphant, T.E., Haberland, M., Reddy, T., Cournapeau, D., Burovski, E., Peterson, P., Weckesser, W., Bright, J., et al. (2020). SciPy 1.0: fundamental algorithms for scientific computing in Python. Nat Methods 17, 261–272. 10.1038/s41592-019-0686-2.

82. Pedregosa, F., Varoquaux, G., Gramfort, A., Michel, V., Thirion, B., Grisel, O., Blondel, M., Prettenhofer, P., Weiss, R., Dubourg, V., et al. (2011). Scikit-learn: Machine Learning in Python. J. Mach. Learn. Res. 12, 2825–2830.

83. Li, C., Fleck, J.S., Martins-Costa, C., Burkard, T.R., Themann, J., Stuempflen, M., Peer, A.M., Vertesy, A., Littleboy, J.B., Esk, C., et al. (2023). Single-cell brain organoid screening identifies developmental defects in autism. Nature 621, 373–380. 10.1038/s41586-023-06473-y.

84. Sullivan, C., and Kaszynski, A. (2019). PyVista: 3D plotting and mesh analysis through a streamlined interface for the Visualization Toolkit (VTK). Journal of Open Source Software 4. 10.21105/joss.01450.

