## Supplemental Figures 1-9 for "Multiscale Spatial Transcriptomic Atlas of Human Basal Ganglia Cell-Type and Cellular Community Organization"

Supplemental Figure 1

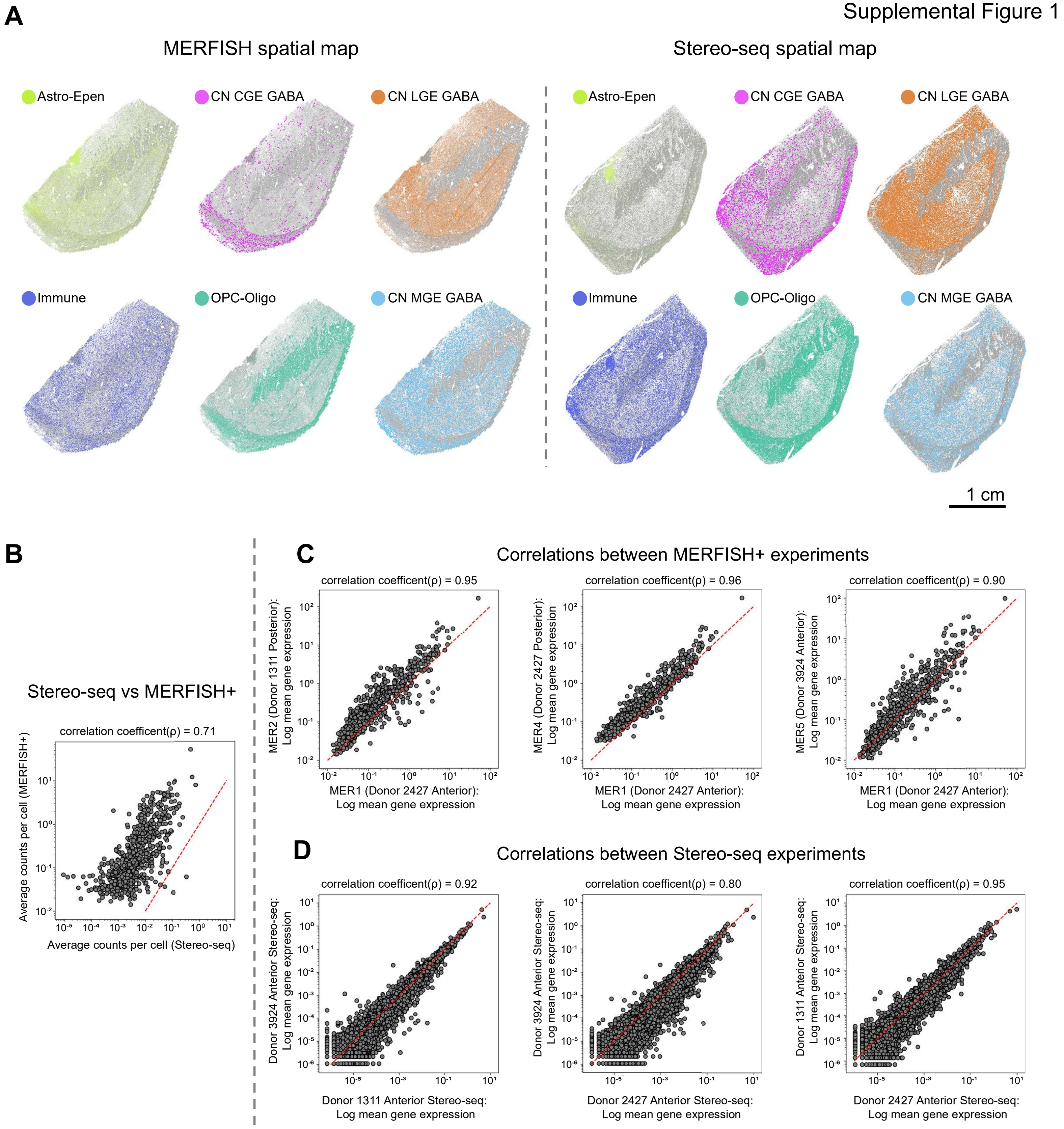

Supplemental Figure 2

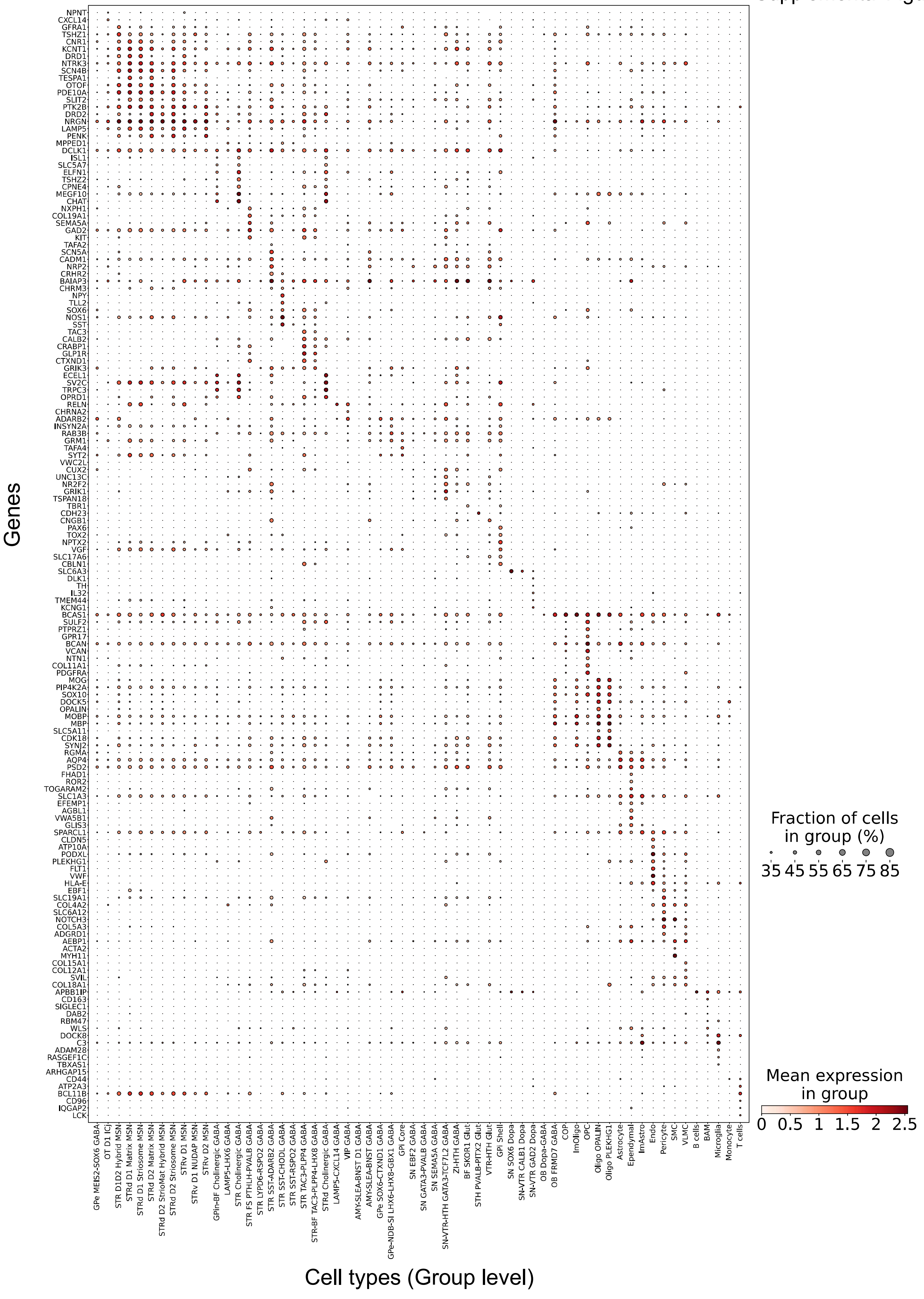

Supplemental Figure 3

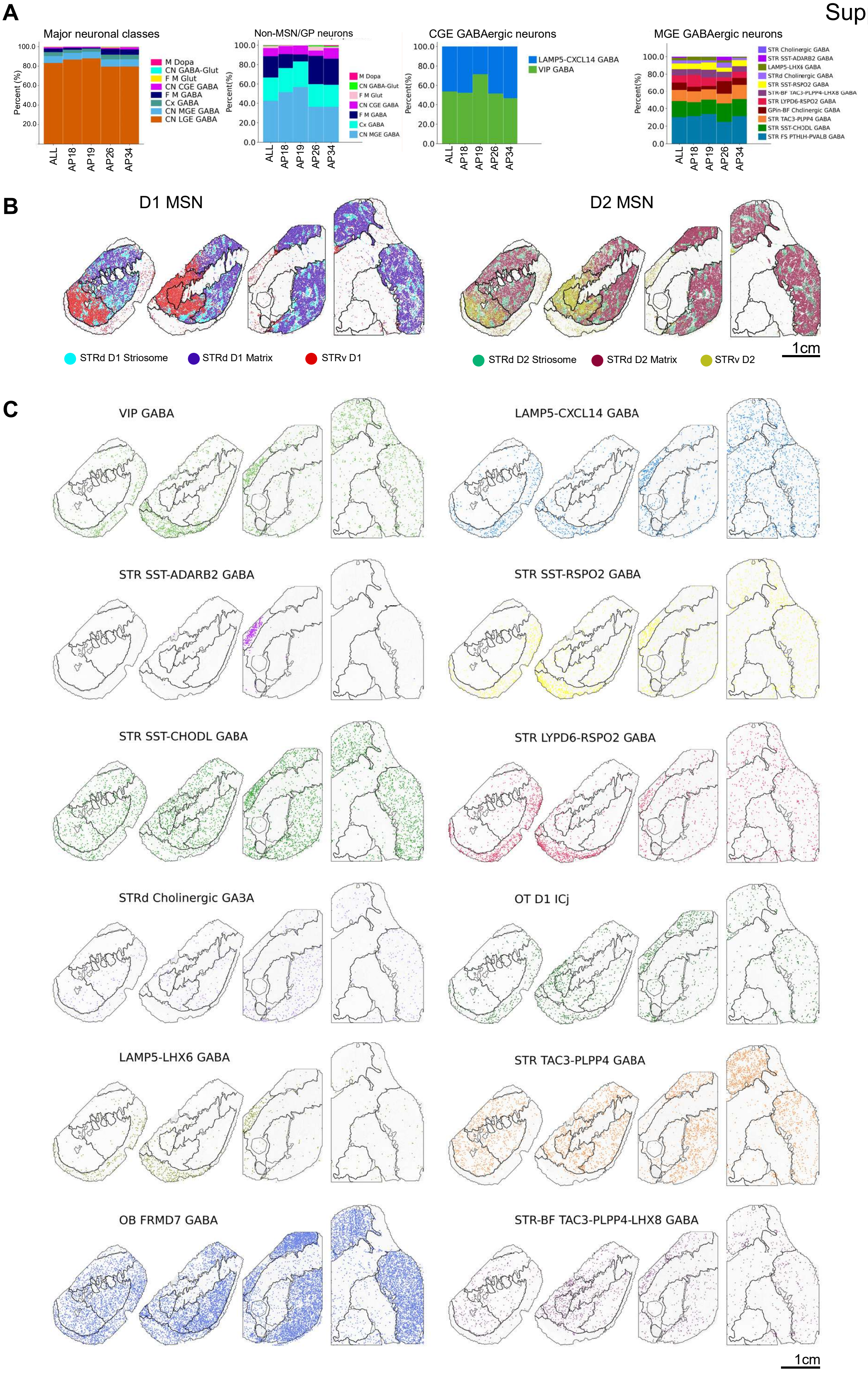

Supplemental Figure 4

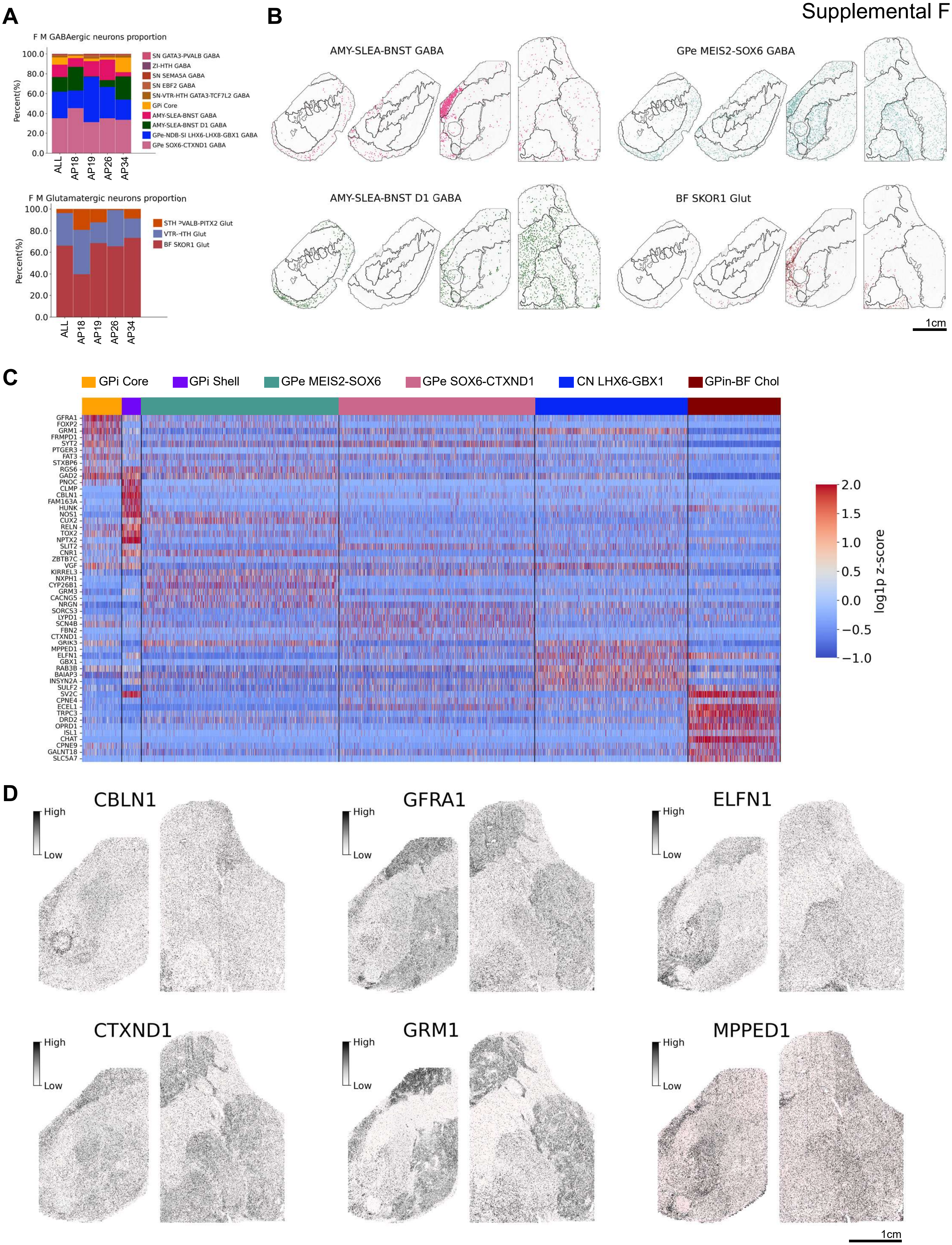

**A**

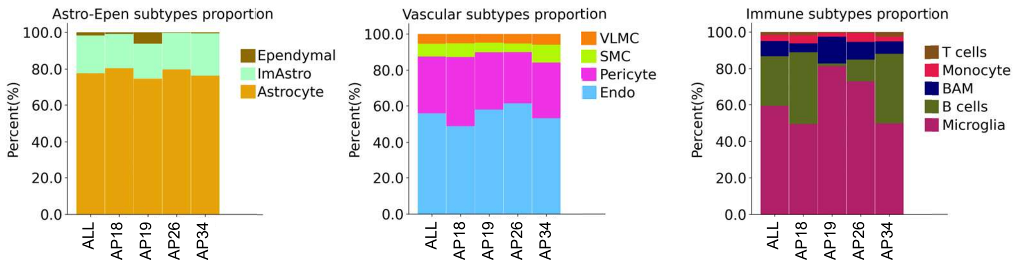

Astro-Epen subtypes proportion
Percent(%)
100.0
80.0
60.0
40.0
20.0
0.0
ALL
AP18
AP19
AP26
AP34
Ependymal
ImAstro
Astrocyte
Vascular subtypes proportion
VLMC
SMC
Pericyte
Endo
Immune subtypes proportion
T cells
Monocyte
BAM
B cells
Microglia


**B**

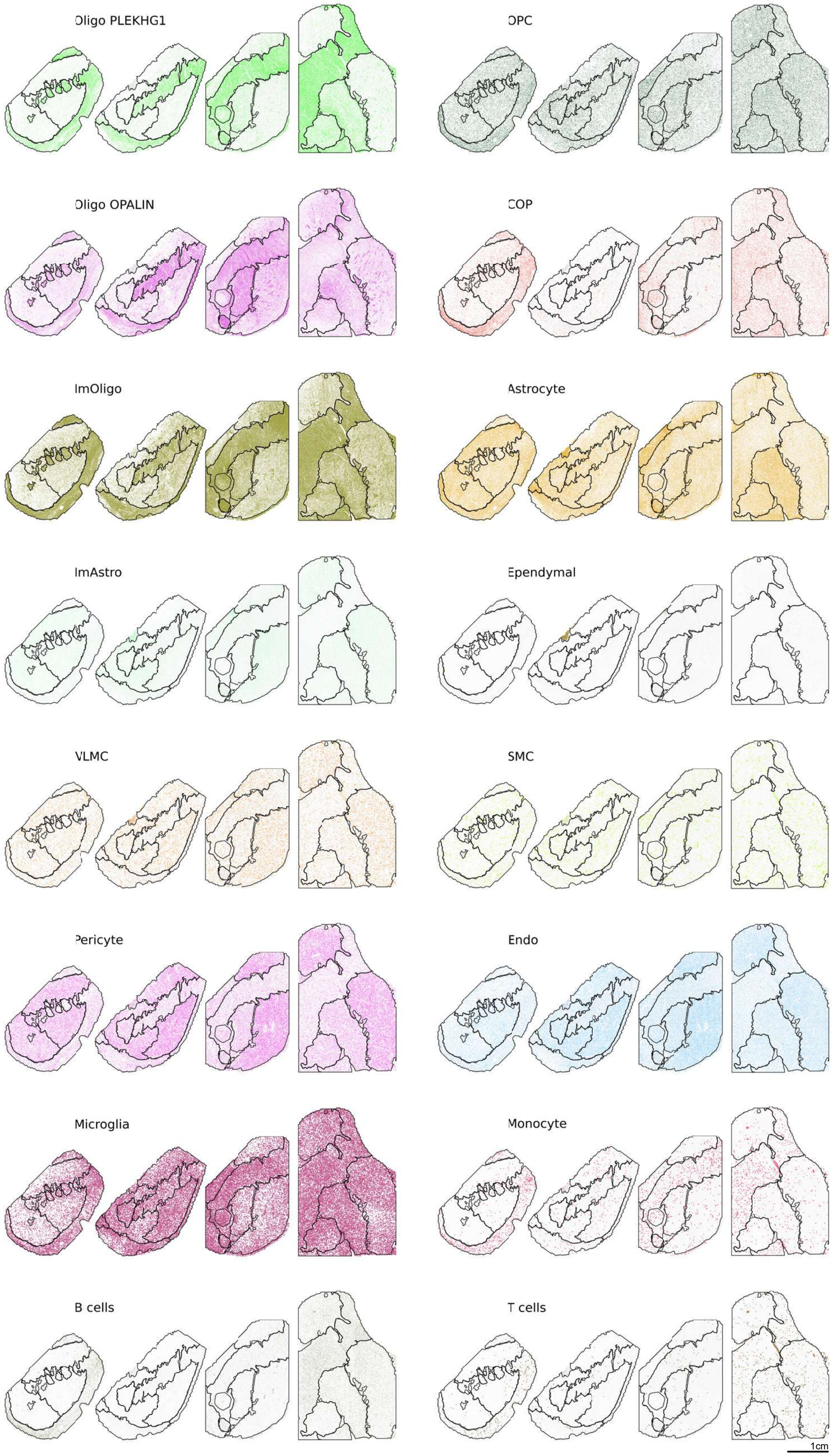

Oligo PLEKHG1
OPC
Oligo OPALIN
COP
ImOligo
Astrocyte
ImAstro
Ependymal
VLMC
SMC
Pericyte
Endo
Microglia
Monocyte
B cells
T cells
1cm

Supplemental Figure 6

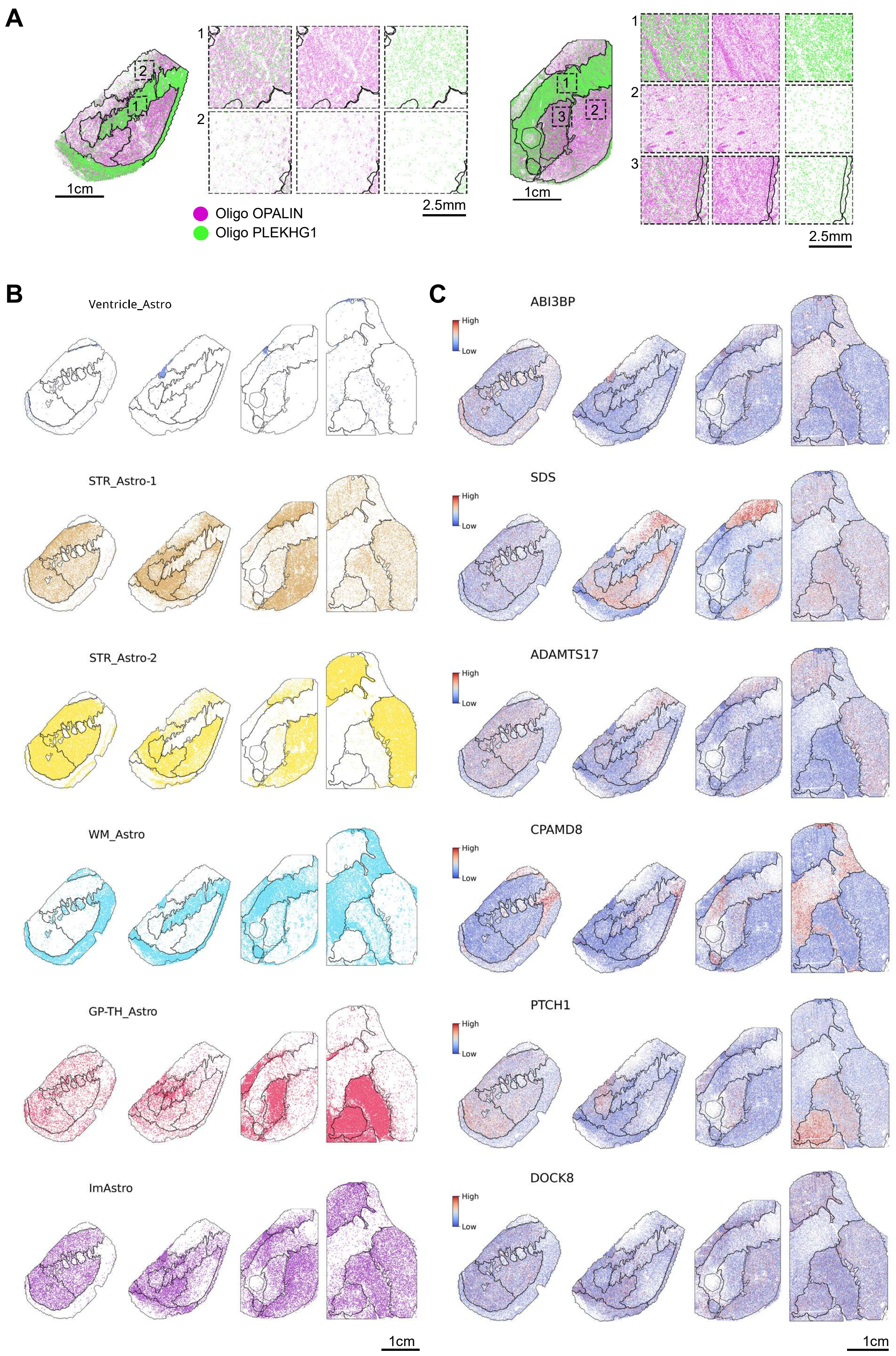

Supplemental Figure 7Su

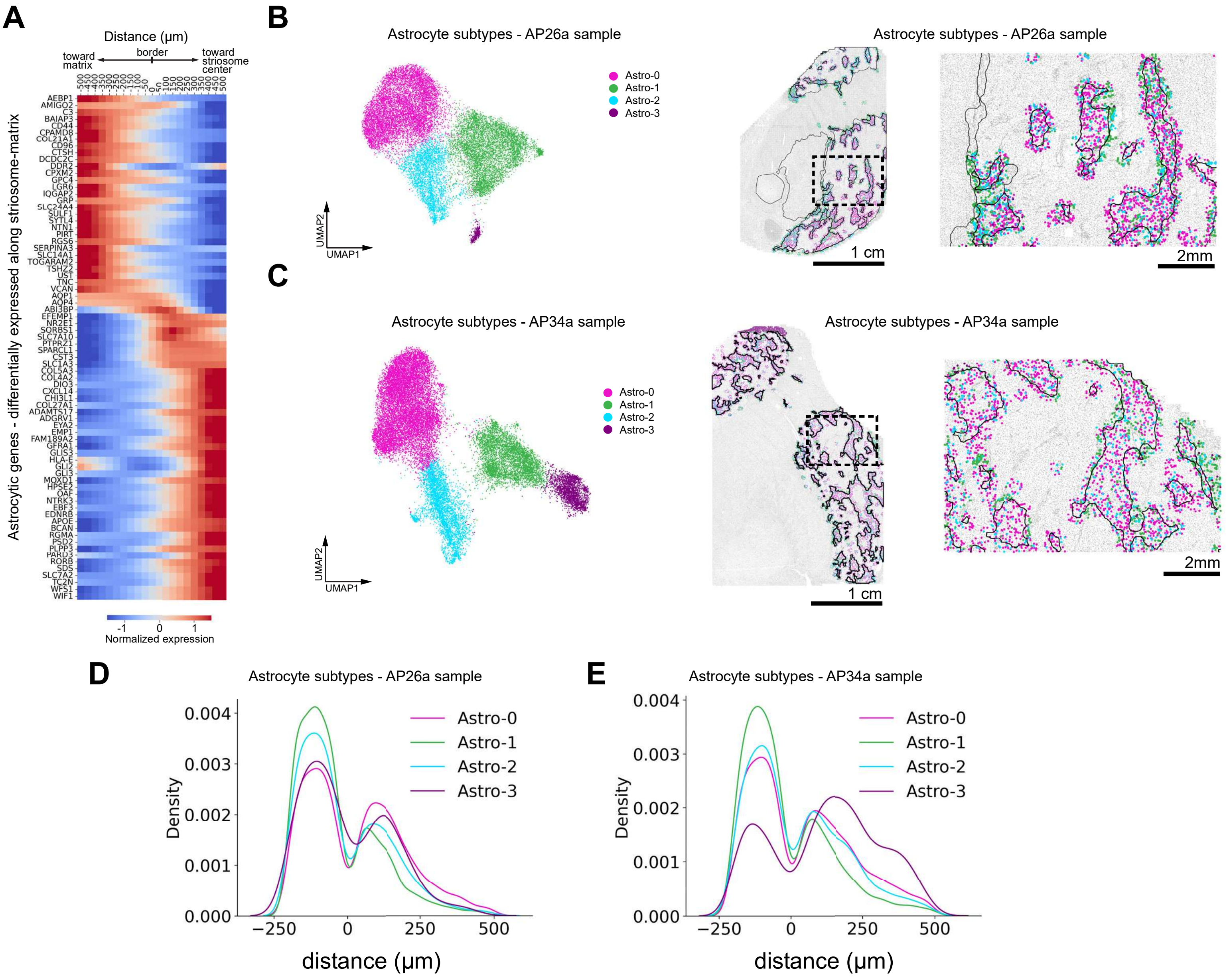

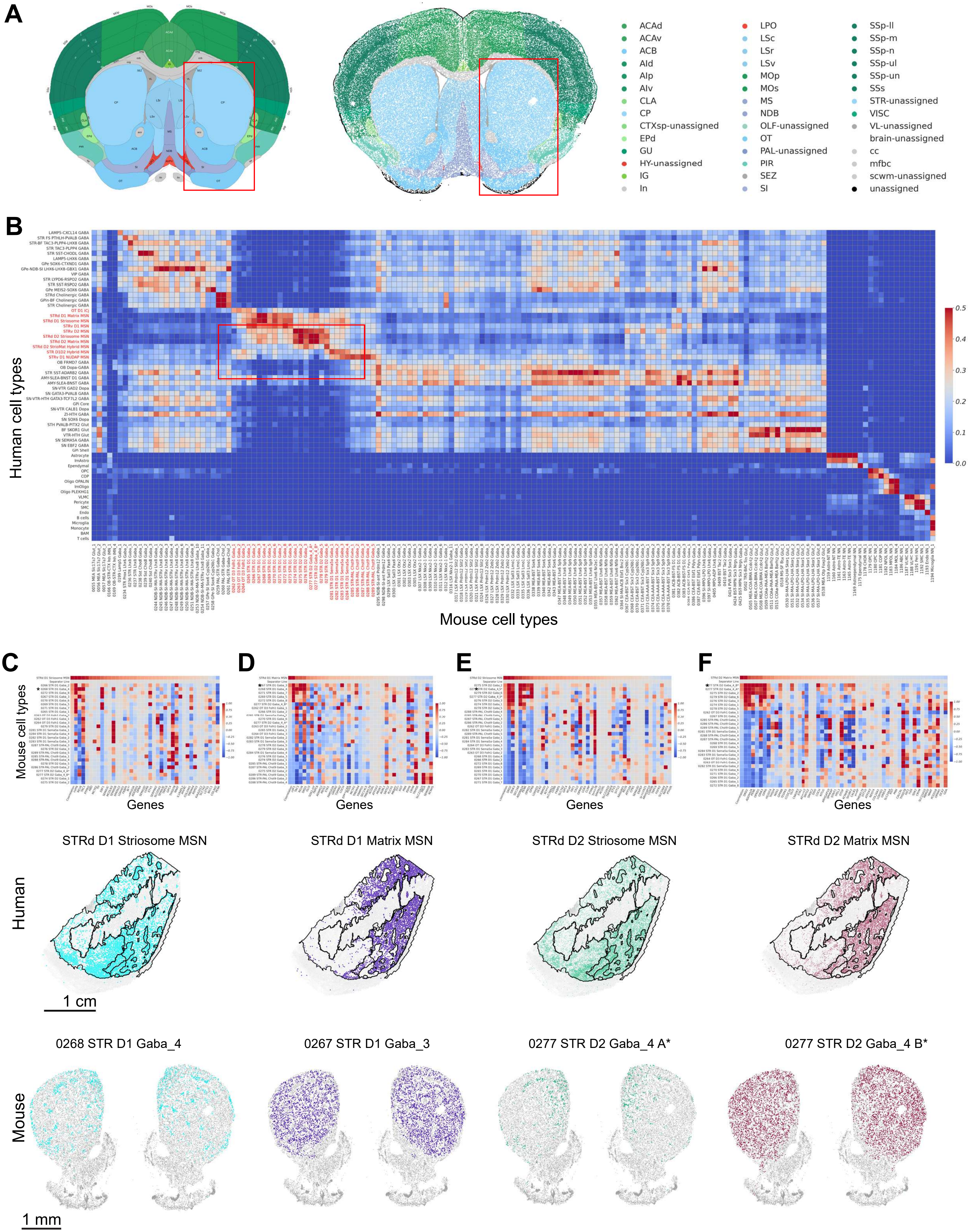


Supplemental Figure 8

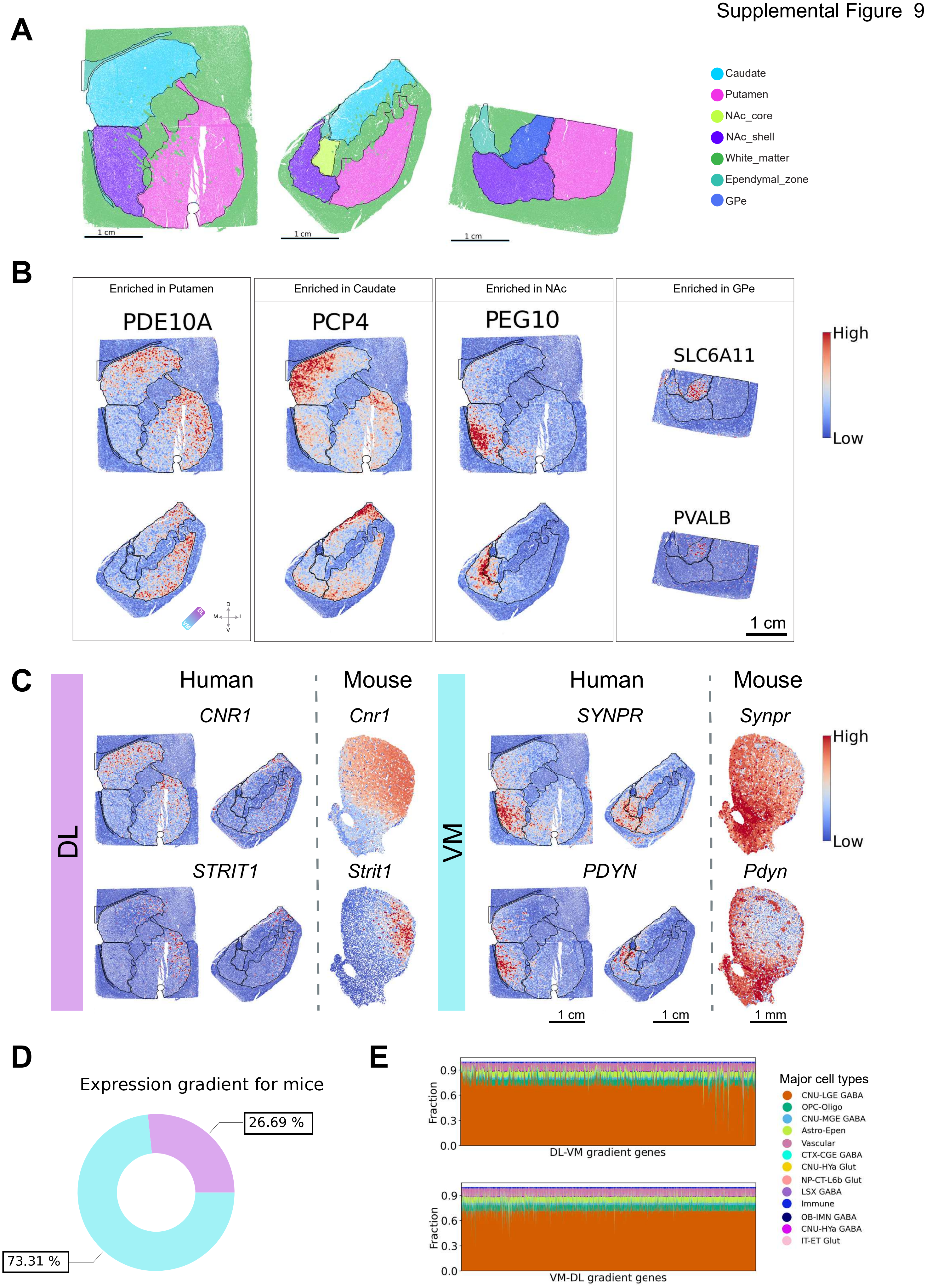
A
Caudate
Putamen
NAc_core
NAc_shell
White_matter
Ependymal_zone
GPe
1 cm
B
Enriched in Putamen
Enriched in Caudate
Enriched in NAc
Enriched in GPe
PDE10A
PCP4
PEG10
SLC6A11
PVALB
High
Low
1 cm
C
DL
VM
Human
Mouse
CNR1
Cnr1
STRIT1
Strit1
SYNPR
Synpr
PDYN
Pdyn
1 cm
1 mm
D
Expression gradient for mice
26.69 %
73.31 %
E
Fraction
DL-VM gradient genes
VM-DL gradient genes
Major cell types
CNU-LGE GABA
OPC-Oligo
CNU-MGE GABA
Astro-Epen
Vascular
CTX-CGE GABA
CNU-HYa Glut
NP-CT-L6b Glut
LSX GABA
Immune
OB-IMN GABA
CNU-HYa GABA
IT-ET Glut
