## Supplemental Table 1 for "Multiscale Spatial Transcriptomic Atlas of Human Basal Ganglia Cell-Type and Cellular Community Organization"

| **Table S1.** Human brain sample information | | | | | | | | |
| --- | --- | --- | --- | --- | --- | --- | --- | --- |
| **Case ID** | **Age** | **Sex** | **Race/Ethnicity** | **PMI (hr)** | **RIN** | **Cause of death** | **Section AP** | **Subregions Profiled** |
| 1311 | 36 | Male | White | 18.3 | 8.2 | Sudden cardiac death associated with severe cardiomegaly | 18a, 34a | Ca, Pu, NAc, IC, GP |
| 5129 | 40 | Male | White | 17.8 | 8.1 | Accidental drowning | 21a | Pu, NAc, IC, GP |
| 3924 | 44 | Male | White/Hispanic | 22.5 | 8.2 | Acute alcohol intoxication | 18a | Ca, Pu, NAc, IC |
| 2724 | 53 | Male | White | 20 | 8.4 | Undetermined (no definitive cause identified) | 19a, 26a | Ca, Pu, NAc, IC, GP |
